# *OsSWEET11b*, a sixth leaf blight susceptibility gene involved in sugar transport-dependent male fertility

**DOI:** 10.1101/2021.08.21.457078

**Authors:** Lin-Bo Wu, Joon-Seob Eom, Reika Isoda, Chenhao Li, Si Nian Char, Dangping Luo, Van Thi Luu, Masayoshi Nakamura, Bing Yang, Wolf B. Frommer

## Abstract

SWEETs play important roles in intercellular sugar transport. Induction of SWEET sugar transporters by transcription activator-like effectors (TALe) of *Xanthomonas* ssp. is a key factor for bacterial leaf blight (BLB) infection of rice, cassava and cotton. Here, we identified the so far unknown OsSWEET11b with roles in male fertility and BLB susceptibility in rice. While single *ossweet11a* or *b* mutants were fertile, double mutants were sterile. Since clade III SWEETs can transport gibberellin (GA), a key hormone for rice spikelet fertility, sterility and BLB susceptibility might be explained by GA transport deficiencies. However, in contrast to the Arabidopsis homologs, OsSWEET11b did not mediate detectable GA transport. Fertility and susceptibility must therefore depend on SWEET11b-mediated sucrose transport. Ectopic induction of *OsSWEET11b* by designer TALe enables TALe-free *Xanthomonas oryzae* pv. *oryzae* (*Xoo*) to cause disease, identifying *OsSWEET11b* as a BLB susceptibility gene and demonstrating that the induction of host sucrose uniporter activity is key to virulence of *Xoo*. Notably, only three of now six clade III SWEETs are targeted by known *Xoo* strains from Asia and Africa. The identification of OsSWEET11b has relevance in the context of fertility and for protecting rice against emerging *Xoo* strains that evolve TALes to exploit *OsSWEET11b*.

## Introduction

Sucrose, produced by photosynthesis, is translocated to other organs such as roots and flowers that depend on carbohydrates supply from leaves. Adequate supply of sucrose to reproductive organs is critical for fertility and therefore yield. The phloem is responsible for sucrose allocation; the rice phloem sap contains ~600 mM sucrose (Hayashi & Chino, 1990). While phloem loading in rice does not seem to make use of apoplasmic transport steps involving SWEET and SUT plasma membrane sucrose transporters, seed filling depends on SWEET and SUT sucrose transporters. Pollen are symplasmically isolated from parental tissues, do not perform photosynthesis, and thus microspores have to be supplied with sugars via the tapetum on an apoplasmic route using plasma membrane sugar transporters. Temporally controlled sugar transport and metabolism likely play roles at multiple locations in the anthers, and dual routes, involving sucrose and hexose transporters, as well as apoplasmic invertases, are required for efficient delivery of sugars to the developing pollen grains. In rice, a cell wall invertase, *OSINV4*, was found to be transiently expressed in the tapetum, and at later developmental stages also in the microspores (Oliver *et al*., 2005). In Arabidopsis, the hexose uniporter AtSWEET8 (originally named Rpg1, Ruptured Pollen grain 1) appears to be involved in the secretion of sugars from the tapetum and together with the hexose/H^+^ symporter AtSTP2 in subsequent uptake into the developing pollen (Truernit *et al*., 1999; Guan *et al*., 2008; Chen *et al*., 2010). Defects in *atsweet8*/*rpg1* mutants include male sterility and defects in primexine, a transient sporophytic carbohydrate layer deposited on the microspore plasma membrane. In addition, the sucrose uniporter AtSWEET14 (also named RPG2, Ruptured Pollen Grain2) also plays a role in pollen nutrition. Later it was found that AtSWEET13 and 14 are expressed in stamina; the exact cellular localization is however not known (Kanno *et al*., 2016); anther dehiscence was delayed in *sweet13*; *sweet14* mutants (Kanno *et al*., 2016). Anther levels of the gibberellins (GA) GA_4_ were ~10x lower in *sweet13*; *sweet14* double mutants relative to wild type Col-0. Notably, GA_3_ was able to supplement the fertility defects. GA is well known to play a role in fertility in both rice and Arabidopsis, in particular, exogenous application of GA can rescue low temperature-triggered sterility (Sakata *et al*., 2014; Kwon & Paek, 2016; Kanno *et al*., 2016). Surprisingly, a yeast three-hybrid (Y3H) screen had identified clade III SWEETs including AtSWEET13 and 14 as GA transporters, raising the possibility that the male sterility in double mutants was caused by either a defect in GA transport, or an indirect effect of a defect in sucrose transport on GA levels (Kanno *et al*., 2016). More recently, the clade I SWEET3a glucose transporter has been shown to be involved in GA transport (Morii *et al*., 2020). In rice, RNA interference of *SWEET11* (Os8N3) also caused male sterility in rice (Yang *et al*., 2006). Surprisingly, however, *sweet11a knock out* mutants generated by CRISPR-Cas9 had no obvious effect on fertility (Yang *et al*., 2018).

SWEETs are well known to mediate cell-to-cell transport of sugars at several key locations in the plant. SWEETs can be grouped into four separate phylogenetic clades, with clade III SWEETs capable of transporting sucrose. Phenotypes of Arabidopsis and maize mutants are consistent with key roles in sucrose transport in many important places, including nectaries, phloem loading and seed filling (Chen *et al*., 2010, 2012; Lin *et al*., 2014; Kanno *et al*., 2016; Bezrutczyk *et al*., 2018, 2021). In rice, OsSWEET11 (from here onwards named OsSWEET11a) is involved in seed filling, particularly in field conditions (Ma *et al*., 2017; Yang *et al*., 2018). OsSWEET14 and OsSWEET15 has overlapping roles with OsSWEET11a (Yang *et al*., 2018; Fei *et al*., 2021). These findings are consistent with roles in sucrose, but it can not be excluded that GA transport plays an important role. Only a dissection of sucrose and GA transport activities using SWEET mutants will enable to determine the relative role of the two activities.

Clade III SWEETs are susceptibility factors for bacterial leaf blight in rice (BLB) (Chen *et al*., 2010, 2012; Eom *et al*., 2019; Oliva *et al*., 2019). Transcription of *OsSWEET11a*, *13* and *14* is directly induced by a suite of TAL effectors (TALes), eukaryotic transcription factors made by the causative bacteria *Xanthomonas oryzae* pv. *oryzae* (*Xoo*). In contrast to five Clade III SWEETs, other hexose transporting members of this family (clade I, II and IV) cannot cause susceptibility when induced by artificial TALes (Streubel *et al*., 2013). It has been hypothesized that the xylem vessel-dwelling bacteria require sucrose as a nutrient, which is not present at sufficient levels in the xylem sap (Bezrutczyk *et al*., 2018). In summary, all five known members of clade III SWEETs can serve as susceptibility factors.

Here we identified a sixth member of the clade III SWEET family in newer rice genome annotations that is closely related to OsSWEET11. OsSWEET11 was herein renamed to OsSWEET11a. The new member is the closest homolog of OsSWEET11a and was named OsSWEET11b. OsSWEET11b mediates transport of sucrose, while transport of GA_3_ and GA_4_ was undetectable. OsSWEET11b levels were highest in anthers, but in contrast to *OsSWEET11a*, not in developing seeds. Patterns of OsSWEET11a and 11b protein accumulation in stamina were complementary. Double *knock out ossweet11a*;*11b* mutants were male sterile. In contrast to the Arabidopsis *atsweet13*;*14* double mutants, GA_3_ did not restore fertility of the double mutant. This work thus identified a new potential BLB susceptibility gene that contributes to male fertility in rice by supporting sucrose transport towards the developing pollen.

## Materials and Methods

### Bioinformatic analyses

Protein sequences of the SWEET genes in Arabidopsis, rice and maize were obtained from Aramemnon (http://aramemnon.uni-koeln.de/), Uniport (https://www.uniprot.org/), and MaizeDB (https://www.maizegdb.org/). The transmembrane domains of SWEETs were predicted using Consensus TM alpha helix prediction (AramTmCon) and TMHMM v2.0 (http://www.cbs.dtu.dk/services/TMHMM/). Alignment of the conserved protein sequences was conducted with MAFFT (Katoh *et al*., 2019) with a gap extension penalty of 0.123 and gap opening penalty of 1.53. Phylogeny tree inference was performed using FastME 2.0 (Lefort *et al*., 2015). The maximum-likelihood method with JTT protein substitution matrix was used to generate phylogeny trees. Clade supporting scores were calculated by 1000 bootstrapping replicates.

### Plant materials and growth conditions

Most experiments were performed using *Oryza sativa* L., ssp. *japonica cv*. Kitaake (Kitaake); infection experiments were in addition performed with *Oryza sativa* L., ssp. *indica cv*. Guanglu’ai 4 (GLA). *Knock out* mutants and translational GUS-fusion lines were in Kitaake background. Dehusked rice seeds were surface-sterilized with 70% ethanol and Klorix^®^bleach solution followed by thorough rinses with autoclaved deionized water. Seeds were placed onto ½ Murashige Skoog media supplemented with 1% sucrose and 0.4% agarose in darkness at 30 °C for 3 days. Emerging seedlings were grown in constant light (200 μmol m^−2^ s^−1^) for one week in a growth chamber. Ten-day-old seedlings were transplanted into 2 L round pots (16.7 cm diameter, 13.2 cm high) filled with soil (Luu *et al*., 2020). Plants were grown in glasshouses at Düsseldorf University (Germany) until maturity; light was supplemented with BL120 LED lamps (Valoya, Finland) to >400 μmol m^−2^ s^−1^, at 30 °C (day) and 25 °C (night), rel. humidity 50-70%. Plants were grown under comparable conditions at Giessen University (Germany). Sterility of the double mutants was also observed at University of Missouri (Columbia, Missouri, USA).

### CRISPR-Cas9 and translational GUS constructs

The single mutant lines *ossweet11a-1*, *ossweet11a-2* and *OsSWEET11a* translational GUS-fusion lines had previously been described (Yang *et al*., 2018). The rice CRISPR system used for creation of *ossweet11b*, and *ossweet11a*;*11b* lines was used as previously described (Zhou *et al*., 2014). Briefly, guide RNAs were designed to target the third exon of *OsSWEET11b* (corresponding to the second transmembrane helix), or both *OsSWEET11a* (targeting the first exon near start codon) and *11b*. Oligonucleotide-derived double strand fragments for the spacer sequences of guide RNAs were cloned into pENTR-gRNA1, followed by Gateway recombination into the Cas9-expressing vector pCas9-GW (Supporting Information Fig. S1, S2). Cas9/gRNA constructs were introduced into *Agrobacterium* strain EHA105; *O. sativa* cv. Kitaake was transformed as described (Hiei *et al*., 1994). Two guide RNAs were designed to target both *OsSWEET11a* and *OsSWEET11b* simultaneously in Kitaake. Independent mutant lines were designated ossweet11b-1, 11b-2 and 11b-3, and the new *osweet11a* alleles in the double mutants *osweet11a-3* and *osweet11a-4* Table S2). The OsSWEET11b-GUS translational fusion construct was generated by inserting the genomic region of *OsSWEET11b* into the pC3000intC derived promoterless GUSplus vector (Yang *et al*., 2018). The genomic fragment of *OsSWEET11b* including a 2155 bp upstream region (calculated upstream of ATG), the complete coding sequence (with introns but without stop codon) was PCR-amplified (primers: Table S1) from genomic DNA of Nipponbare. PCR products were purified and cloned into the GUSplus vector digested with *XbaI*/*BamHI* restriction enzymes. Fusion constructs were sequenced for validation and introduced into the wild type Kitaake (Fig. S3). Thirty-one transformants were obtained while seven showed comparable GUS activity in stamen. Two lines, 2-7-2 and 9-2-2 of T_3_ generation were used for in detail characterization.

### Genotyping of CRISPR plants

Leaf genomic DNA was isolated using a peqGOLD Plant DNA Mini Kit (PeqGold, VWR International GmbH, Darmstadt, Germany) following the manufacturer’s instructions. To identify CRISPR/Cas9-mediated mutations, PCR was performed using High-Fidelity Phusion PCR Master Mix (NEB, Frankfurt am Main, Germany) (initial denaturation (98 °C, 30 s), 30 cycles of reaction (98 °C, 10 s; 66/60 °C for *OsSWEET11a*/*11b*, 15 s; 72 °C, 30 s) and final extension (72°C, 2 mins) (primers: Table S1)). PCR amplicons were purified using NucleoSpin Gel and PCR Clean-up kits (Macherey-Nagel, Düren, Germany) and subjected to Sanger sequencing for identifying mutations. Chromatograms were analyzed with SnapGene Viewer (CSL Biotech LLC, San Diego, USA). DNA sequences were aligned using MEGA-X software (Kumar *et al*., 2018). Genotyping results were summarized in Fig. S4 and Table S2. The new *ossweet11a* mutant alleles were designated as *ossweet11a-3* and *ossweet11a-4* (Fig. S4, Table S2).

### Phenotypic analyses of mutant plants

Wild type Kitaake, two independent lines of *ossweet11a and ossweet11b* single and *ossweet11a*;*11b* double mutants were grown until maturity. To examine sterility, rice spikelets were collected 1 d prior to anthesis and fixed in 70% ethanol at room temperature. After removing lemma and palea with dissecting forceps, whole florets and carpels were observed and documented under a Axiozoom V.16 (Carl Zeiss, Jena, Germany) stereo zoom microscope. For analyses of pollen grains, anthers were removed from the florets and dissected on glass slides for Lugol’s KI-I_2_ staining (Sigma). Grain development at different stages was recorded using a stereo microscope camera (Nikon DS-Fi3) after dissection in two-day intervals (2-10 days after flowering) by careful dehusking of immature seeds. Mature panicles were photographed using a digital camera (Fujifilm X-T3).

### Reciprocal crosses for sterility investigation

Reciprocal crosses were made to test whether sterility was caused by defects in male or female gametogenesis in *ossweet11a*;*11b* double *knock out* lines. Crosses were performed as follows: *ossweet11a-3*;*11b-3* (♀, progenies of line 1-5-6, Table S2) × wild type (♂); wild type (♀) × *ossweet11a-3*;*11b-3*(♂) *ossweet11a-3*;*11b-3* (♀) × *ossweet11a-2* (♂, (Yang *et al*., 2018)), *ossweet11a-2* (♀) × *ossweet11a-3*;*11b-3* (♂), *ossweet11a-3*;*11b-3* (♀) × *ossweet11b-1* (♂, line 9-1-4, Table S2); *and ossweet11b-1* (♀) × *ossweet11a-3*;*11b-3* (♂). Anthers of recipient parent spikelets were carefully removed with sharp scissors 2-3 d before anthesis. Emasculated spikelets were covered with paper bags until artificial pollination performed between 11 am and 1 pm. Panicles from donor plants at heading stage were cut for artificial pollination. Pollinated spikelets were kept in paper bags until maturity. F_1_ seeds from crosses were photographed using a stereo microscope camera (Nikon DS-Fi3).

### Sugar transport assays using FRET sucrose sensors

ORFs of rice clade III SWEETs (*OsSWEET11a*, *11b*, *12*, *13*, *14*, and *15*) were cloned by PCR into the Gateway^™^ entry vector pDONR221f1 (Chen *et al*., 2012; Yang *et al*., 2018). LR reactions were performed to transfer ORFs into the mammalian expression vector pcDNA3.2V5. Sanger sequencing was employed to verify all constructs. Transport assays were performed in 96-well plates (Chen *et al*., 2010). HEK293T cells were co-transfected with a plasmid carrying the sucrose sensor FLIPsuc90μ-sCsA (Sadoine *et al*., 2021) and a plasmid carrying a candidate transporter gene (100 ng) using Lipofectamine2000 (Invitrogen). For FRET imaging, culture media in each well were replaced with 100 μl Hanks Balanced Saline Salt (HBSS) buffer followed by addition of 100 μL HBSS buffer containing 25 mM sucrose. A Leica inverted fluorescence microscope DM IRE2 with Quant EM camera was used for imaging with SlideBook 4.2 (Intelligent Imaging Innovations) with exposure time 200 msec, gain 3, binning 2, and time interval 10 sec (Hou *et al*., 2011).

### GA transport assays in mammalian cells

For gibberellic acid (GA) transport assays, the ORF of *OsSWEET11b* was amplified (Table S1) and assembled with a mammalian expression vector pcDNA3.1 Hyg(+) digested with *BamHI* and *XhoI* using NEBuilder HiFi DNA Assembly Master Mix (New England Biolabs). HEK293T cells were co-transfected with constructs carrying the GA sensor GPS1 (Rizza *et al*., 2017) and *OsSWEET11b* by Lipofectamine LTX (Invitrogen) in 8-well glass bottom chambers (Iwaki Cat#; 5232-008) and incubated for 48 h. Culture medium was replaced with 300 μL Dulbecco’s Modified Eagle Medium (D-MEM) without phenol red containing either 0.001% (v/v) DMSO or 1.0 μM GA_3_ dissolved in 0.001% DMSO and cells were incubated for 3 h. Fluorescence was acquired by a Nikon Ti2-E microscope equipped with 40x lens under excitation at 440 nm with two emission channels for CFP and YFP. Images were taken for 3 min with 5-sec intervals. Image quantification was performed with Fiji/ImageJ software (NIH).

### GA transport assays in a Yeast-3-Hybrid system

The ORF of *OsSWEET11b* was cloned into the yeast expression vector pDRf1-GW which contains a strong *PMA1* promoter fragment to drive high levels of expression in yeast cells (Loqué *et al*., 2007) by LR Gateway recombination. GA transport assays were performed using a previously described Y3H system (generous gift of Mitsunori Seo, RIKEN, Yokohama)(Kanno *et al*., 2016). The yeast strain PJ69-4a [MATa *trp1-901 leu2-3*,*112 ura3-52 his3-200 Δga14 Δga180* LYS2::GAL1-HIS3 GAL2-ADE2 *met2*::GAL7-lacZ] was co-transformed with the GA receptor components pDEST22-GAI and pDEST32-GID1a and either pDRf1-GW (empty vector control), pDRf1-AtSWEET13, or pDRf1-OsSWEET11b by conventional lithium acetate/PEG transformation. Three independent colonies were used as technical replicates for each assay. Colonies were inoculated in synthetic defined (SD -Leu, -Trp, -Ura) liquid media and incubated overnight at 30°C. the culture was diluted sequentially to 10, 10^2^, 10^3^, and 10^4^ cells/μL. 10 μL cell suspension was spotted on SD (-Leu, -Trp, -Ura) or selective (-Leu, -Trp, -Ura, -His) media containing 3 mM 3-amino-1,2,4-triazole (3-AT) and 0.001% (v/v) DMSO, and 0.1 μM GA_3_ or 1 nM GA_4_ in 0.001% (v/v) DMSO, and incubated for 3 days at 30°C. Plates were photographed.

### Exogenous application of GA on rice plants

Plants of *ossweet11a*;*11b* double mutants in R2 stage (collar formation on flag leaf) were sprayed with 10 μM GA_3_ dissolved in 0.1% (v/v) DMSO as a foliar application. Foliar spray of 20 ml 10 μM GA_3_ solution was applied repeatedly for three days between 10:00 and 11:00 each day (mock: 0.1% DMSO. Impact of GA_3_ application on shoots was documented 7 d after last foliar spray using a digital camera (Fujifilm XT-3). Florets of *ossweet11a;11b* double mutants in both mock and GA_3_ treatment were dissected and observed under a stereo zoom microscope Axiozoom V.16 (Zeiss, Germany). Mature pollen was stained with Lugol’s KI/I_2_ solution and documented on an Axiozoom V.16 stereo zoom microscope.

### Subcellular localization in *Nicotiana benthamiana* leaves

To amplify the coding region of *OsSWEET11b*, total RNA was isolated from young spikelets of Kitaake (RNeasy Plant Mini Kit, Qiagen) and cDNA was synthetized (Maxima^™^ H Minus cDNA Synthesis Master Mix, Thermo Fisher Scientific). The coding sequence of *OsSWEET11b* was PCR amplified without STOP codon (primers: Table S1) using Phusion High-Fidelity PCR polymerase (Thermo Fisher Scientific) and cloned into the Gateway donor vector pDONR221 (Thermo Fisher Scientific). The entry vector harboring *OsSWEET11b* was then included in an LR reaction (Thermo Fisher Scientific) with pAB117 (provided by Prof. Dr. Rüdiger Simon, HHU Düsseldorf) that contains the β-Estradiol-inducible CaMV 35S promoter and eGFP coding sequence. The final expression plasmid pAB117:SWEET11b:eGFP carrying *OsSWEET11b* fused at the C-terminus with the enhanced green fluorescent protein (eGFP) and driven by the β-estradiol-inducible CaMV 35S promoter was generated and validated by Sanger sequencing. *Agrobacterium tumefaciens* GV3101 was transformed with pAB117:SWEET11b:eGFP. *Agrobacterium* culture preparation and tobacco leaf infiltration were performed as described (Sosso *et al*., 2015). As a control, pAB118:AtMAZZA:mCherry carrying the coding sequence of *Arabidopsis* MAZZA gene fused at the C-terminus with mCherry driven by the β-estradiol inducible CaMV 35S promoter was used (Blümke *et al*., 2021). Fluorescence was detected on an Olympus SpinSR with excitation/emission 488/522–572 nm (eGFP) and 5 and 561/667-773 nm (chlorophyll). Epidermal leaf chloroplast fluorescence was used to differentiate vacuolar or cytosolic localization (lining chloroplasts on the vacuolar side) or plasma membrane localization (peripheral to chloroplasts). Image analysis was performed using Fiji (https://fiji.sc/). Experiments were repeated twice using 2-5 different *Agrobacterium* colonies and 4-10 *N. benthamiana* plants per construct.

### Histochemical GUS activity analysis and paraffin sectioning

Lines harboring pOsSWEET11a:OsSWEET11a-GUS and pOsSWEET11b:OsSWEET11b-GUS translational fusions were analyzed as described (Yang *et al*., 2018). In brief, rice seedlings and spikelets at the mature pollen stage were harvested and pre-fixed in 90% ice-cold acetone by 10 min vacuum infiltration. After 30 min incubation at room temperature, spikelets were transferred to GUS washing buffer (50 mM sodium phosphate buffer pH 7.0, 1 mM potassium ferrocyanide, 1 mM potassium ferricyanide, 10 mM EDTA, 0.1% (v/v) Triton X-100 and 20% (v/v) methanol) with vacuum infiltration for 10 min on ice. After removal of washing buffer, samples were vacuum infiltrated in GUS staining solution (GUS washing buffer containing 2 mM 5-bromo-4-chloro-3-indolyl-beta-D-glucuronide, X-gluc) for 10 min in the dark. Specimen were incubated for 2 hours at 37°C in darkness followed by an ethanol series (20%, 30%, 50% and 70%), 30 minutes each, at room temperature. Observation and documentation of GUS activity were performed under a stereo microscope Axiozoom V.16. For paraffin sections, X-gluc-stained specimen were fixed in FAA solution containing 50% (v/v) ethanol, 3.7% (v/v) formaldehyde and 5% (v/v) acetic acid for 30 min at room temperature. After removing the fixative, samples were dehydrated with an ethanol series (30 min each; 80%, 90%, 95% and 100%) and 100% tert-butanol. Histoplast paraffin (Leica Biosystems, Nussloch, Germany) was melted at 60 °C for sample embedding. Sections (10 μm) were cut with a rotary microtome and mounted on SuperFrost Plus slides (Fisher Scientific, Schwerte, Germany). Sectioned samples were observed with a light microscope (CKX53, Olympus, Hamburg, Germany) and documented with an EP50 camera (Olympus).

### Synthesis of designer TALe gene and disease assays

Designer dTALe for *OsSWEET11b* was assembled from a library of 51 individual repeats as described (Li *et al*., 2013). Briefly, the modular repeats with the repeat variable di-residues (RVDs) at positions 12 and 13 (i.e., NI, HD, NG and NN that recognize A, C, T, and G, respectively) were ligated into an array of octamer repeats in the pTLN vector using Golden Gate assembly. Three arrays of octamers were digested using *Sph*1 and *Pst*I for the first octamer, *Pst*I and *BsrG*I for the second octamer, and *BsrG*I and *Aat*II for the third octamer. Recovered fragments were ligated into pZW-ccdB-dTALe predigested with *Sph*I and *Aat*II, resulting in a designer TALe gene corresponding to 24 nucleotides of the target site. The dTALe gene was mobilized into a pHM1-based vector pHM1-Gib compatible with *Xoo* and *E. coli* (Li *et al*., 2019). The resulting pHM1-dTALe was introduced via electroporation into ME2, a *pthXo1*-inactivated mutant of PXO99^A^ (Yang & White, 2004). Bacteria (OD_600_ 0.25) were used for infection by leaf tip clipping; lesion lengths as indicators of virulence were measured as described (Yang & White, 2004). Bacterial infiltration with inoculum (OD_600_ 0.5) was performed for gene expression analyses by qRT-PCR using 3-week-old Kitaake plants. Data were analyzed using one-way ANOVA and Tukey honest significant difference for post-AVOVA pair-wise tests for significance, set at 5% (*p*<0.05).

## Results

### *SWEET11b*, a sixth clade III SWEET gene in the rice genome

A BLASTp search for SWEET homologs in the updated rice genome annotation (Phytozome, *Oryza sativa Kitaake v3.1*; *Oryza sativa Japonica Group Annotation Release 102*) led to the identification of a new member of the SWEET gene family on chromosome 9 (*Gramene*: *Os09g0508250*; NCBI Reference Sequence: XP_015611383.1). The closest paralog of this gene is *OsSWEET11a* (LOC_Os08g42350; Os8N3; Alphafold AF-Q6YZF3-F1) with 64% identity to *SWEET11b* (Fig. S5). The exon-intron structure of the two genes is conserved (Fig. S6a). The genes are located on different chromosomes and thus are not the result of a recent tandem duplication. Phylogenetically SWEET11b falls into clade III (Fig. 1). Notably, a new annotation of the maize genome (B73 RefGen_V3) also contains a new SWEET11 homolog, which is most closely related to OsSWEET11b, and was thus named ZmSWEET11b (Fig. 1).

**Figure 1.**
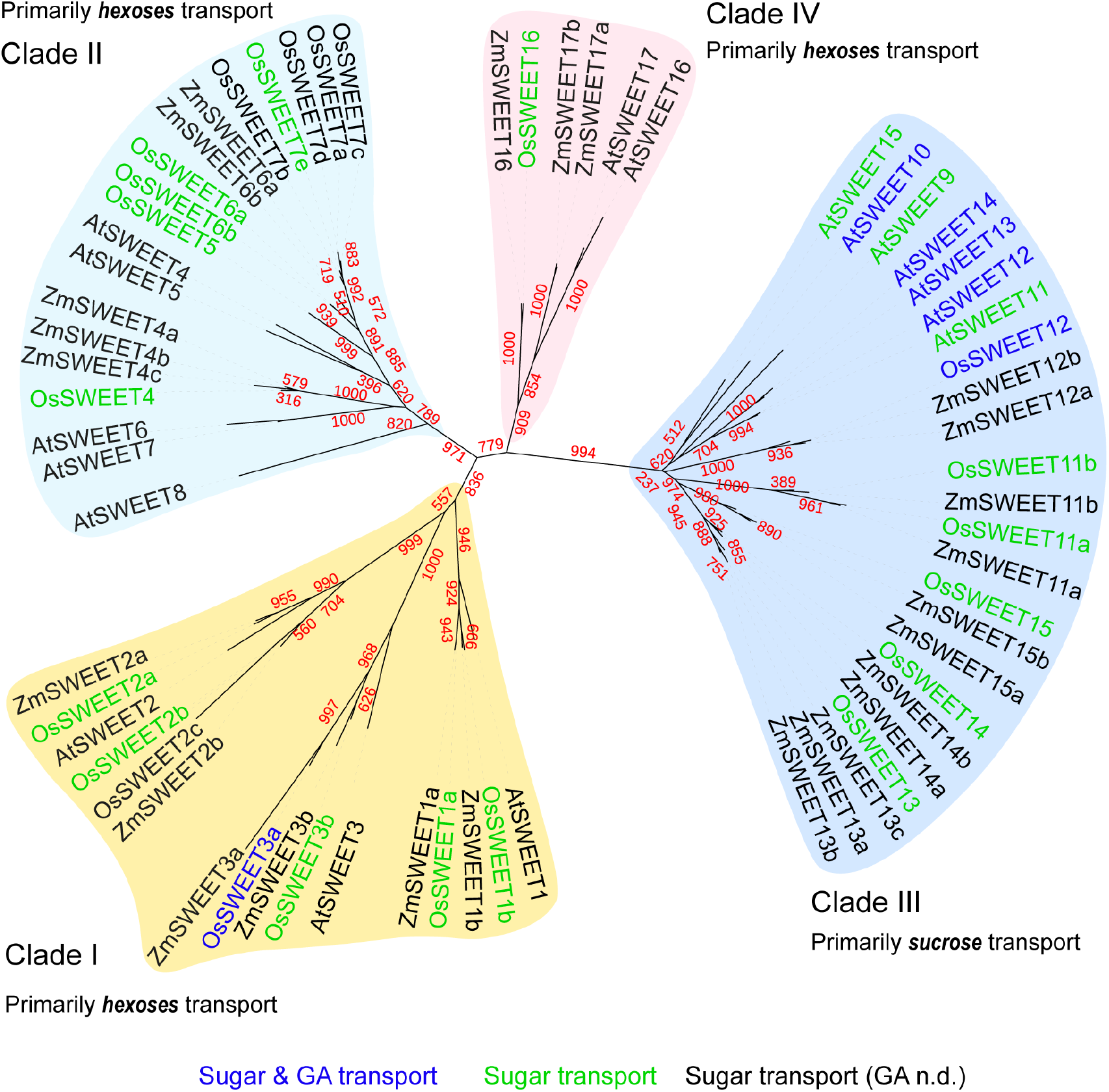
Phylogeny of SWEET gene family members in *Arabidopsis* (At), rice (Os) and maize (Zm). Unrooted phylogenetic tree was generated based on the conserved transmembrane regions of the protein sequences with the help of NGphylogeny (https://ngphylogeny.fr/) and visualized with iTOL program (https://itol.embl.de/). Clade clustering scores (in red) shown as the bootstrap replicates were calculated based on bootstrapping (n = 1000). The SWEETs were highlighted based on the sucrose/GA transport capability. Blue color indicates the SWEETs capable of both sucrose/hexoses and GA transport while green and purple colors show the SWEETs only capable of sucrose/hexoses transport or GA transport activity has not been determined (Kanno *et al*., 2016; Morii *et al*., 2020). GA, gibberellic acid; n.d., not determined.

### Complementary roles of OsSWEET11a and b in anthers

*OsSWEET11a* has a key role in seed filling (Ma *et al*., 2017; Yang *et al*., 2018; Fei *et al*., 2021). Based on public RNAseq data (NCBI BioProject: PRJNA243371), *OsSWEET11a* and *11b* are both expressed in florets as well, and *OsSWEET11b* transcripts accumulate in developing seeds, however only to low levels (Wang *et al*., 2015)(Fig. S7). To assess whether the two paralogs are redundant and have overlapping cell-type specificity, the localization of OsSWEET11a and 11b was investigated using translational GUS fusions driven by their own promoters (2155 bp for *OsSWEET11b*, Fig. S3; 2106 bp for *OsSWEET11a* (Yang *et al*., 2018)). GUS activity from both OsSWEET11a and b translational fusions was detected in stamina and the veins of lemma and palea (Fig. 2a, b). However, in the stamina, the patterns were non-overlapping and GUS activity was neither detected in the tapetum nor in microspores. OsSWEET11a-derived GUS activity was detected in the tip of the filament, i.e., the anther peduncle (Fig. 2c, e). OsSWEET11b expression was detected in the veins of the anther starting in the region where OsSWEET11a activity terminated (Fig. 2c, d). After dehiscence, OsSWEET11b was also found in the vascular bundle of anthers (Fig. 2d, e). OsSWEET11a thus appears to play a role in release of substrates from the vasculature in the basal zone of the anther, while OsSWEET11b likely functions in anther veins directed towards the developing microspore. In addition to SWEET11b protein accumulation in florets, SWEET11b-derived GUS activity was also detected in the stele of the primary roots, emerging lateral roots, leaf veins and spikelet branches (Fig. S8).

**Figure 2:**
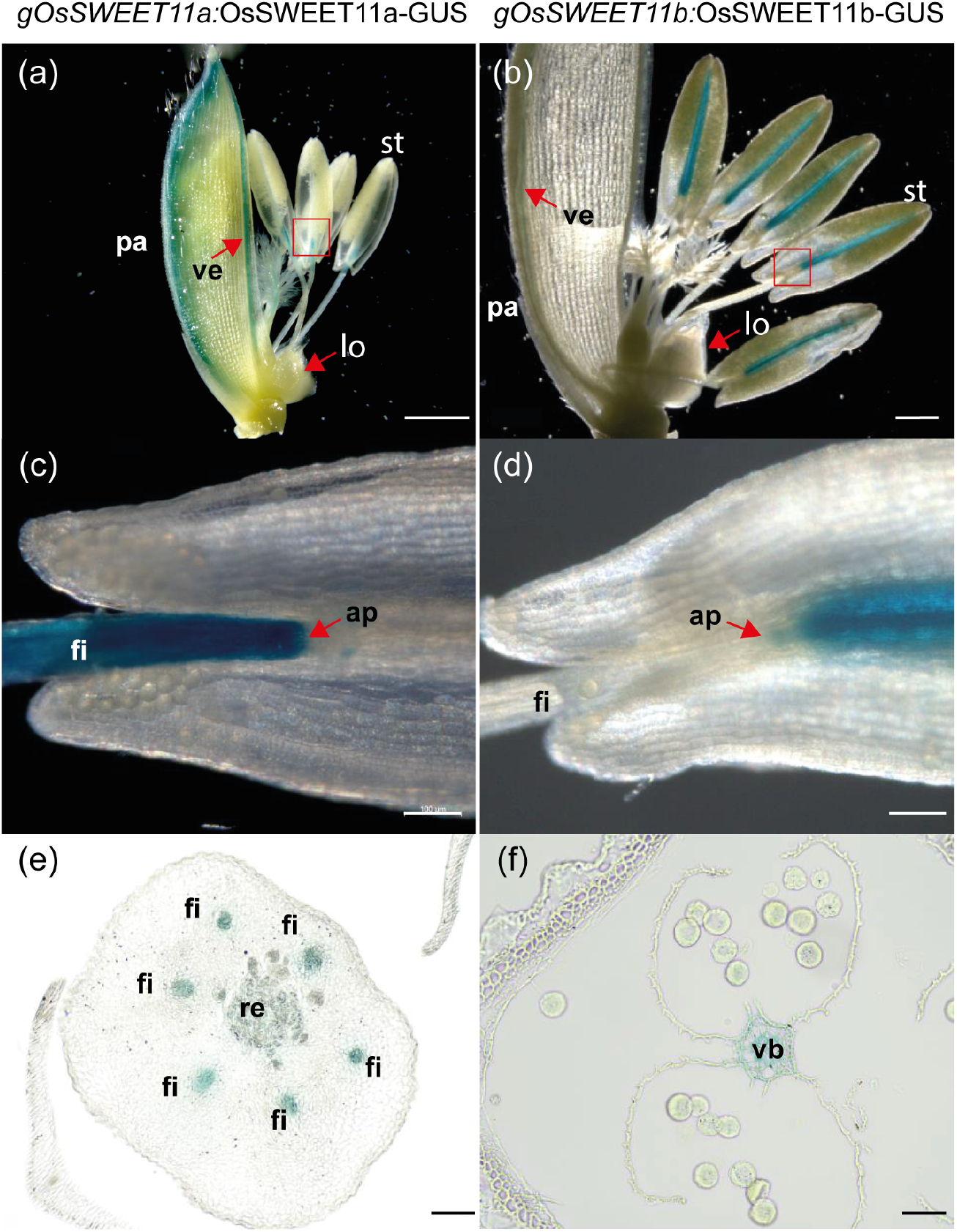
Cell type specific accumulation patterns for OsSWEET11a and OsSWEET11b using GUS histochemistry in florets. (a) Floret at mature pollen stage of *gOsSWEET11a*:OsSWEET11a-GUS line #10 with lemma removed. GUS activity was detected in veins (ve) and filament (fl). (b) Dissected floret at mature pollen stage of *gOsSWEET11b*:OsSWEET11b-GUS line #2. GUS activity was detected in veins and vascular bundle (vb) of anthers. (c) Area in (a) indicated by red square, showing OsSWEET11a-derived GUS activity in the filament that ends in the anther peduncle (ap). (d) Part of stamen (st) is indicated by red square in (b). GUS activity from OsSWEET11b-GUS starts above the anther peduncle and is limited to the vascular bundle of anthers. (e) Transverse section of lower floret in *gOsSWEET11a*:OsSWEET11a-GUS line, showing GUS activity in filament and receptacle (re). (f) Transverse section of upper floret in *gOsSWEET11b*:OsSWEET11b-GUS line, showing GUS activity in vascular bundle of anthers. Scale bars: (a) 1000 μm; (b) 500 μm; (c, d, e, f) 100 μm. Comparable results with eighteen and seven independent transgenic lines for OsSWEET11a and 11b, respectively.

### OsSWEET11b functions as a plasma membrane sucrose transporter

Due to the high similarity of OsSWEET11b to the sucrose transporting OsSWEET11a, one may expect that OsSWEET11b has similar properties, i.e., plasma membrane localization and sucrose transport activity. Confocal imaging of a translational OsSWEET11b-GFP fusion in *N. benthamiana* confirmed plasma membrane localization (Fig. S9, S10). Coexpression of OsSWEET11b with the Förster Resonance Energy sucrose sensor FLIPsuc-90μΔ1V in mammalian HEK293T cells demonstrated that similar to its paralogs, OsSWEET11b mediates sucrose transport (Fig. 3, Fig. S11).

**Figure 3.**
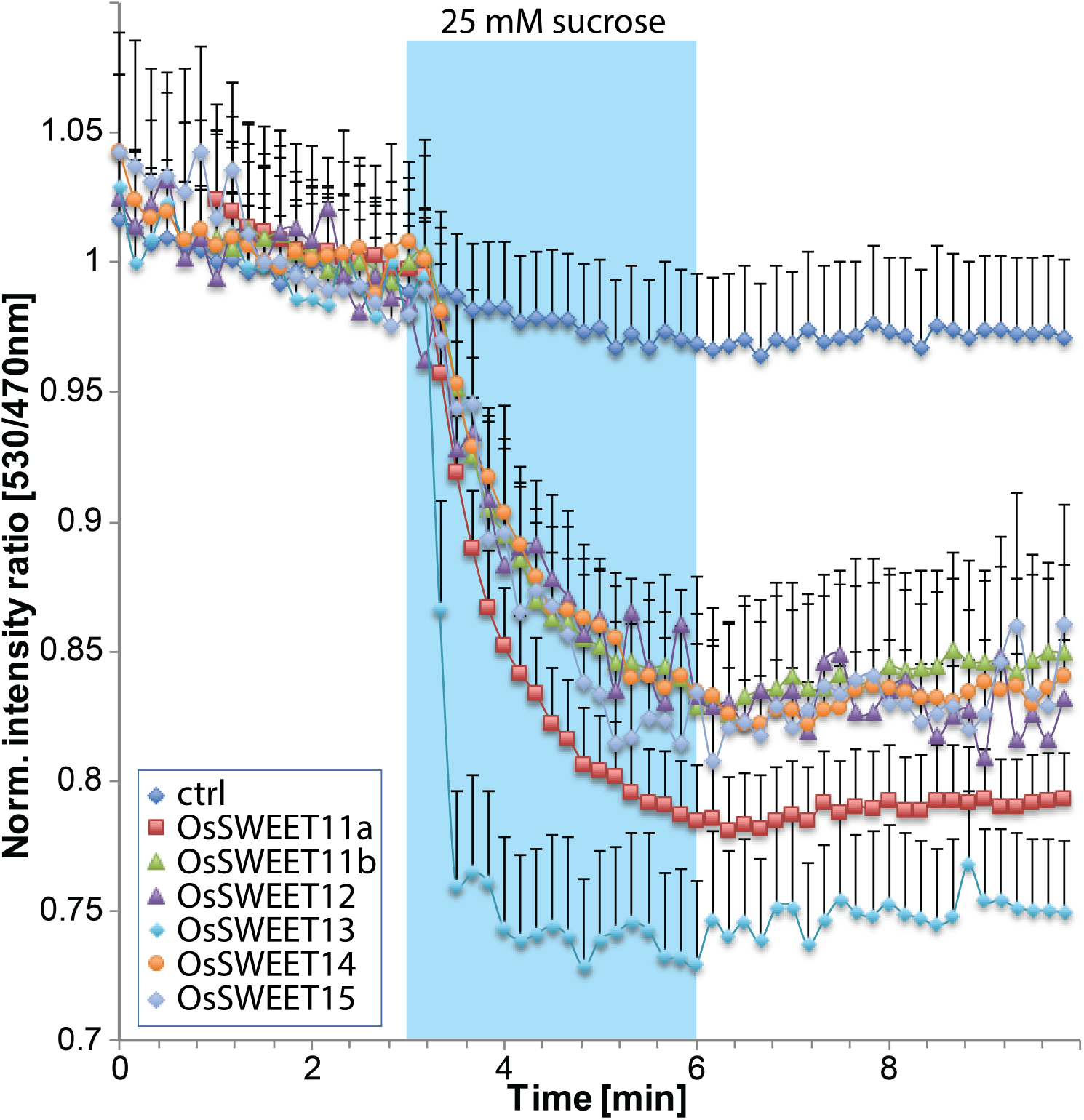
Sucrose transport activity of OsSWEET11b in HEK293T cells. All six *OsSWEETs* in clade III were coexpressed with the sucrose sensor FLIPsuc90μ. Perfusion with a square pulse of 25 mM sucrose led to a negative ratio change, consistent with the accumulation of sucrose in HEK293T cells. In the absence of SWEETs (ctrl, vector only), no significant change in ratio was observed (mean + S.E.M; n= 8 cells doe OsSWEET11b; others between n=5 and n=13). Experiment was repeated at least three times independently.

### Male sterility of *sweet11a*,*b* double *knock out* mutants

To determine the physiological role of OsSWEET11b, *knock out* mutants were generated using CRISPR/Cas9 with a guide RNA targeting the second transmembrane domain (Table S2, Fig. S4, S5c). Panicles of single *knock out* mutant lines *ossweet11b-1* and -*2*, both carrying frameshift mutations (1-bp insertion; 1-bp deletion and one SNP) resulting in a predicted new polypeptide that contained only the first 31 amino acids (aa) of OsSWEET11b (293 aa), showed no obvious phenotypic differences to the Kitaake controls, and florets were fertile when grown in three different locations and several seasons (greenhouses in University of Missouri, Düsseldorf or Giessen) (Fig. 4, Fig. S12a). In addition, seed filling appeared normal (Fig. 4b, Fig. S12b). As described before, *ossweet11a-1*, and -*2 knock out* mutants (Yang *et al*., 2018), grown in parallel, did not show fertility defects. The *stay green* phenotype of the panicles of *ossweet11a* mutants previously observed was not observed here in greenhouses at Düsseldorf and Giessen (Fig. 4a, Fig S12). In contrast, two independent alleles of the double mutant *ossweet11a*;*11b* showed a *stay green* phenotype at Giessen and Düsseldorf, and were sterile (Fig. 4a, b, Fig. S12). Alleles with in-frame mutations causing single amino acid deletions did not lead to sterility (Table S2). Analysis of seed development indicated that defects were likely due to male or female infertility (Fig. 4c). Under Düsseldorf greenhouse conditions, the seed filling defect of *ossweet11a-1* and -*2* was not detectable (Fig. S12b). The phenotypes for the same genotypes are likely conditional; the severity of the seed filling phenotype of *ossweet11a* mutants was previously shown to be more severe under field conditions (Ma *et al*., 2017). Based on the presence of both OsSWEET11a and b proteins in stamina, one may hypothesize that the sterility is due to defects in male gametogenesis. Consistent with this hypothesis, female organs appeared normal (Fig. S13). To test for the source of fertility defects more directly, reciprocal crosses were performed. Results demonstrate that under the same growth conditions, double mutants were male sterile and that double mutant pollen was defective and unable to produce a single fertile seed (Fig. 5a, Fig. S14). Notably, defects were also visible as a waxy appearance of anthers (Fig. 5b, Fig. S15d). Pollen of double mutants frequently had abnormal shapes, and starch staining indicated that the pollen was inadequately supplied with carbohydrates since starch content was substantially lower compared to wild type pollen (Fig. 5c). Thus, based on accumulation in the anther peduncle (OsSWEET11a) and anther veins (OsSWEET11b), a combined defect in the transfer of SWEET substrates from these two locations causes insufficient translocation to the pollen, resulting in a reduction in starch content and thus male sterility. The phenotype could either be a direct effect of a sucrose transport deficiency in anther veins, or a reduced supply with the key hormone GA or its precursors.

**Figure 4.**
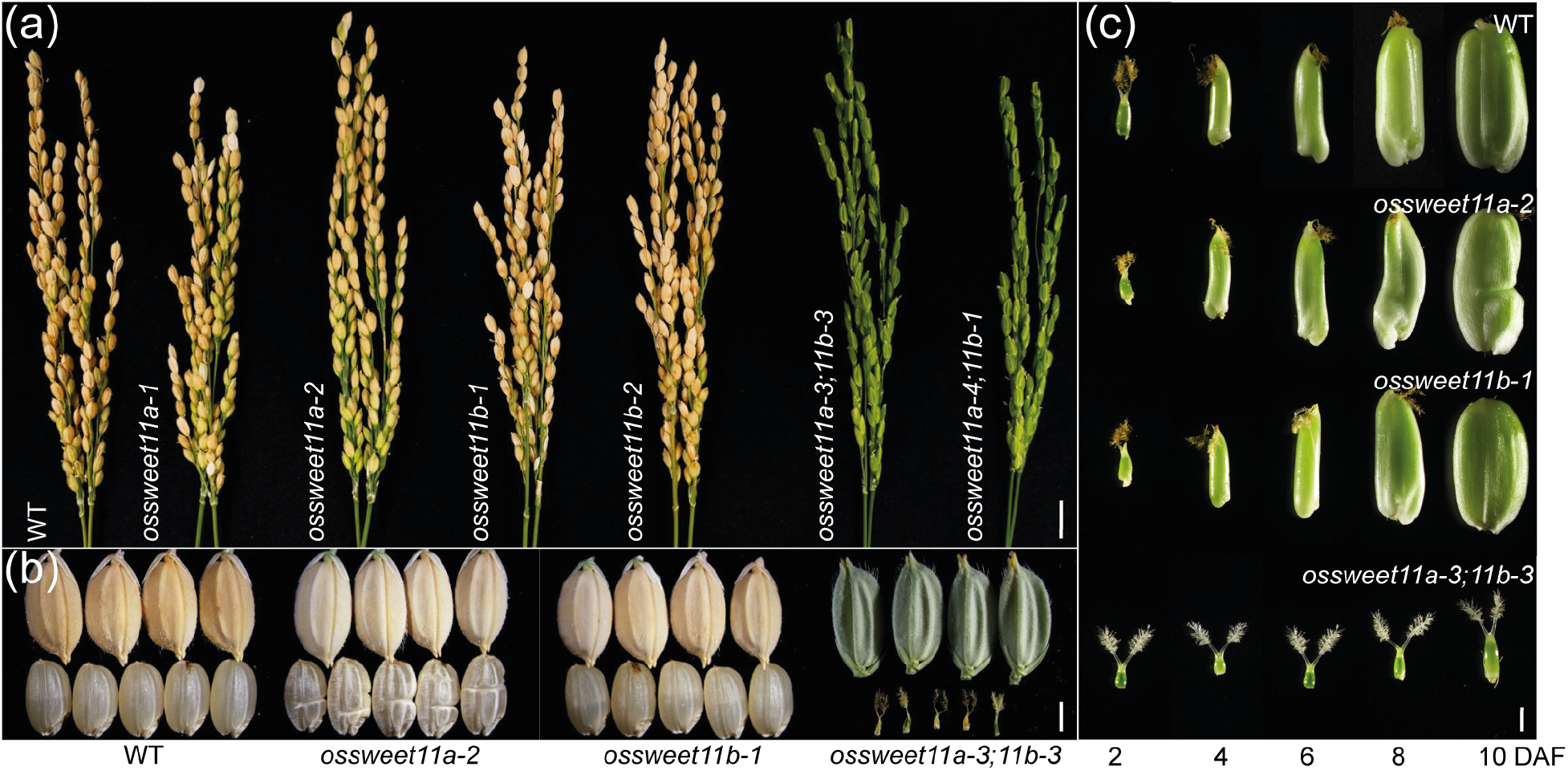
Phenotypes of mature panicles and seed development of wild type Kitaake, *ossweet11a*, *ossweet11b* and *ossweet11a*;*11b* double mutants. (a) Mature panicles from wild type Kitaake (WT), *ossweet11a-1*, *ossweet11a-2*, *ossweet11b-1*, *ossweet11b-2* and two independent combinations of *ossweet11a*;*11b* double mutant alleles 40 days after flowering (DAF). (b) Rice grains with (upper) and without (lower) husks in WT, *ossweet11a-2*, *ossweet11b-1*, and *ossweet11a-3*;*11b-3 double mutants. ossweet11a-2* single mutants showed incomplete seed-filling. *ossweet11a-3*;*b-3* double mutant plants did not develop seeds. (c) Grain development at 2, 4, 6, 8 and 10 days after flowering. *ossweet11a-3*;*11b-3* double mutant plants did not develop seeds. Scale bars: (a) 1 cm; (b) 2 mm and (c) 1 mm.

**Figure 5.**
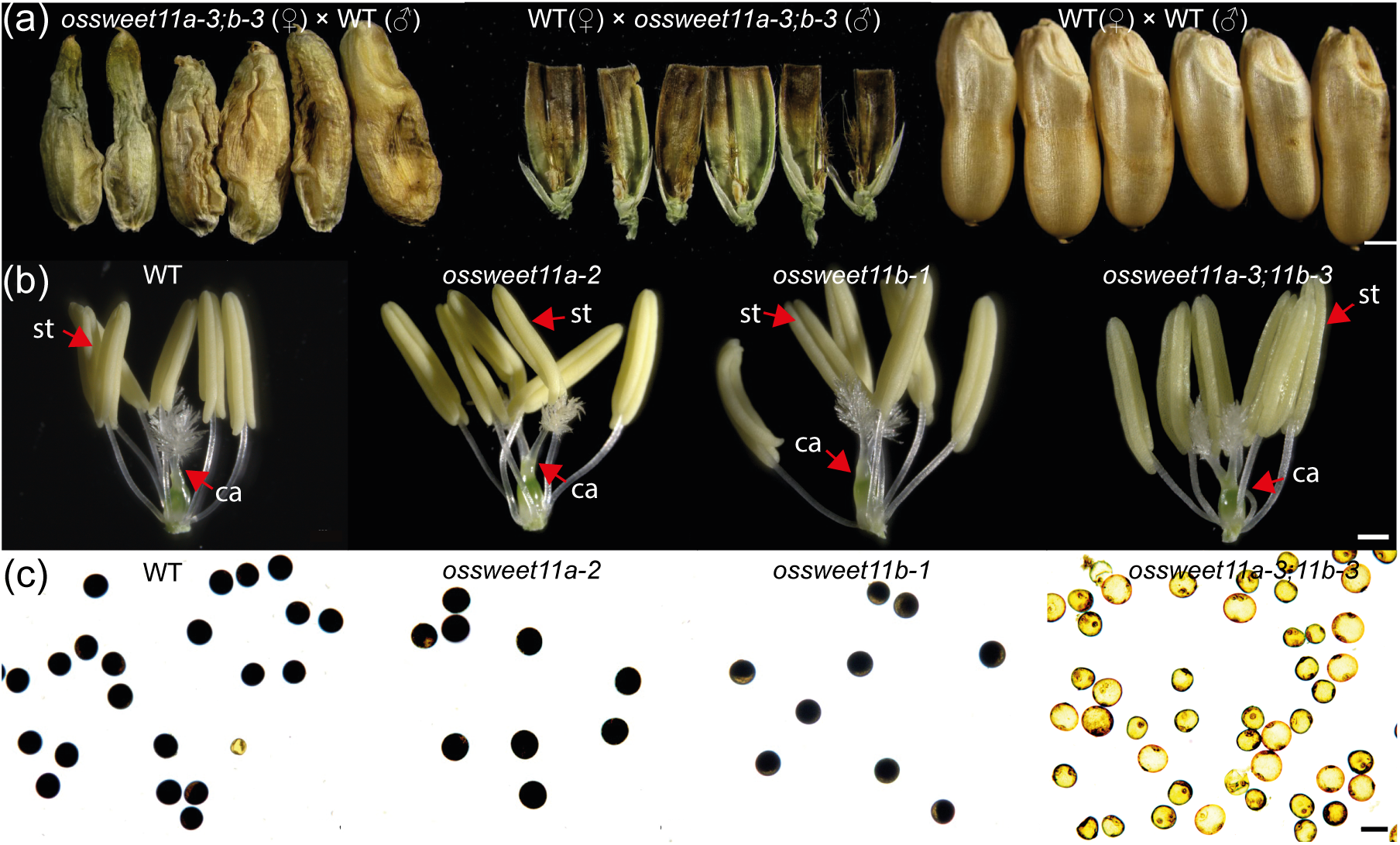
Investigation of the cause of sterility in *ossweet11a*;*11b* double mutant by reciprocal crosses, flowers observation and pollen starch staining. (a) Reciprocal crosses between *ossweet11a-3*;*11b-3* and wild type Kitaake. Wild type Kitaake (WT) as donor parent (♂) restored the fertility in double mutants. F_1_ seeds of double mutant and WT showed incomplete seed filling phenotype. (b) Florets without palea and lemma of *ossweet11a-2*, *ossweet11b-1* and *ossweet11a-3*;*11b-3*. *ossweet11a-3*;*11b-3* had waxy anthers, other genotypes showed normal anther development. (c) Pollen stained with Lugol’s KI/I_2_ solution in WT, *ossweet11a-2*, *ossweet11b-1* and *ossweet11a-3*;*11b-3*. Pollen of double mutant lines was irregular and lacked starch. St: stamen, ca, carpel. Scale bars: (a) 1 mm; (b) 200 μm; (c) 50 μm.

### GA supplementation did not suppress male sterility

GA is required for male fertility in rice. Since the close homologs of Arabidopsis AtSWEET13 and 14 can transport sucrose and GA, and since GA application supplemented male sterility of *atsweet13*;*14* double mutants (Kanno *et al*., 2016), one might speculate that GA could also supplement *ossweet11a*;*11b* fertility defects (Kanno *et al*., 2016). Double mutant plants repeatedly sprayed with a solution containing 10 μM GA_3_ at stage R2 (collar formation on flag leaf) showed a substantial increase in height and internode length, demonstrating that GA entered the plant and had the expected growth promoting effects (Fig. 6a). GA_3_ supplementation had however no positive effect on the fertility of *ossweet11a*;*11b* mutants (Fig. 6b-d). Therefore, it was hypothesized that OsSWEET11b might be able to transport sucrose, but in contrast to the Arabidopsis homologs unable to transport GA. To test this hypothesis, the transport capacity of OsSWEET11b was tested using two independent assays systems – using GPS1, a FRET-based GA sensor in mammalian cells, and a GA-dependent Y3H assay (Kanno *et al*., 2016; Rizza *et al*., 2017). As a control, AtSWEET13 was co-transfected with the GA sensor GPS1 into human HEK293T cells. GPS1 showed significantly higher YFP/CFP fluorescence ratios in the presence of AtSWEET13 supplied with 1 μM GA_3_ relative to controls (Fig. 7, Fig. S16), indicating that GA_3_ was transported by AtSWEET13 into the HEK293T cells. In contrast, OsSWEET11b did not show significant changes in YFP/CFP fluorescence ratio upon GA_3_ treatment, indicating that while sucrose transport is detectable at levels comparable to other SWEETs, OsSWEET11b is unable to mediate measurable GA uptake. As an independent approach, GA uptake assays were performed using a GA-uptake dependent Y3H (Kanno *et al*., 2016). While AtSWEET13 rescued growth on - His media in the presence of GA_3_ or GA_4_, OsSWEET11b, driven by the strong PMA1 promoter fragment, did not enable detectable growth, indicating that OsSWEET11b is not able to provide substantial GA transport activity (Fig. 7, Fig. S17). This result is similar as described in a parallel study for LOC_Os09g32992 (defined here as OsSWEET11b), a gene not characterized further by the authors (Morii *et al*., 2020). Notably, substantial GA transport was also not detected for OsSWEET11a (Morii *et al*., 2020. Thus, in contrast to the Arabidopsis clade III SWEETs, OsSWEET11b is unable of ineffective in mediating GA transport, implying that the sterility observed in *ossweet11a*;*11b* mutants was due to defects in sucrose transport, and not GA supply.

**Figure 6.**
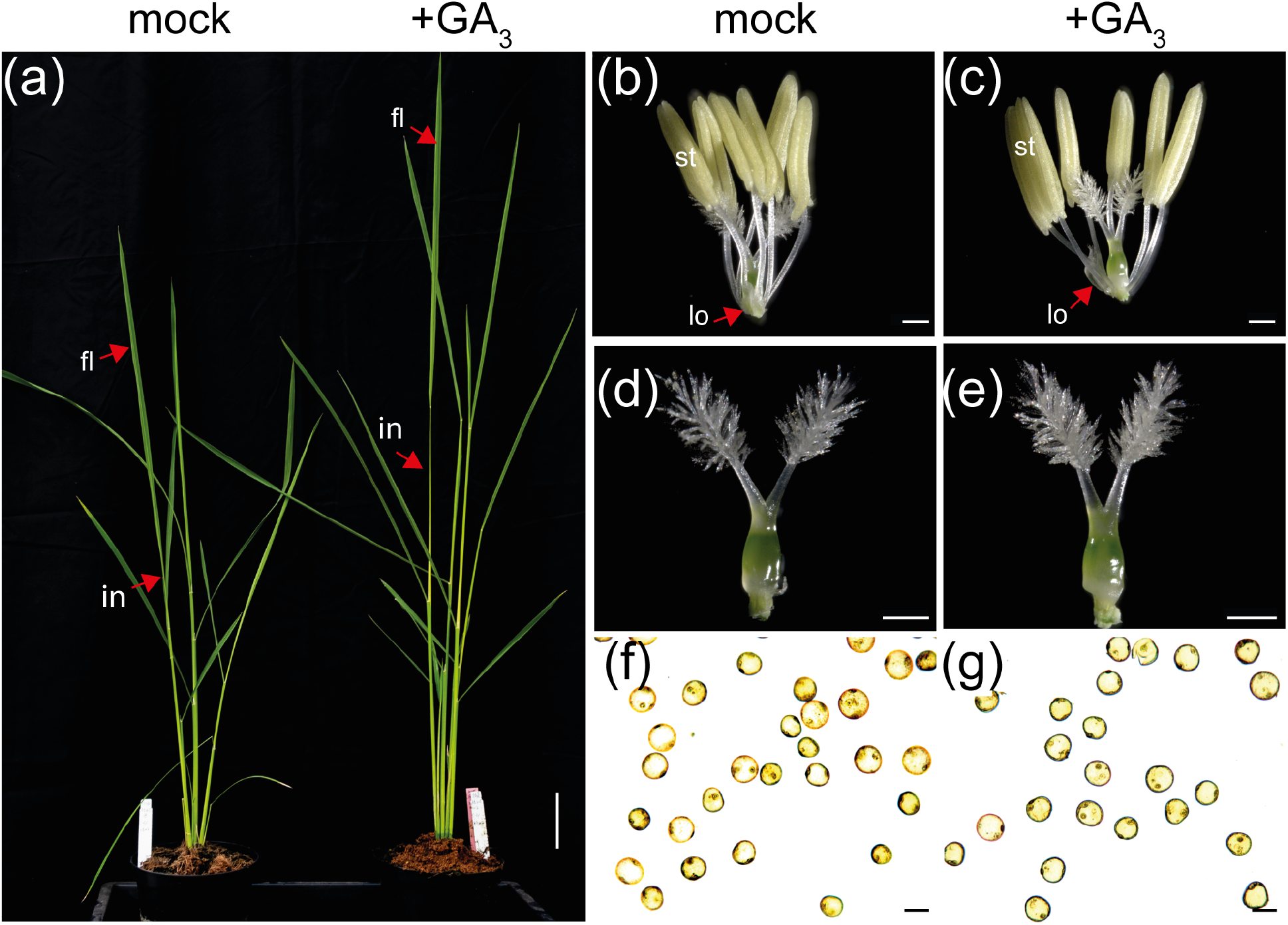
External application of GA did not rescue male fertility defects in *osweet11a*, *b* double mutants. (a) Effect of GA_3_ application on the growth of *sweet11a-3;11b-3* double mutants(left: mock; right: 10 μM GA_3_). Floral organs (b, d) and Lugol’s KI/I_2_ stained pollen (f) of *sweet11a-3*;*11b-3* double mutants without GA_3_ application. Floral organs (c, e) and pollen (g) of *sweet11a-3*;*11b-3* double mutants treated with 10 μM GA_3_. in, internode; fl, flag leaf; lo, lodicule; st, stamen. Scale bars: (a) 10 cm; (b - e) 500 μm; (f, g) 50 μm.

**Figure 7.**
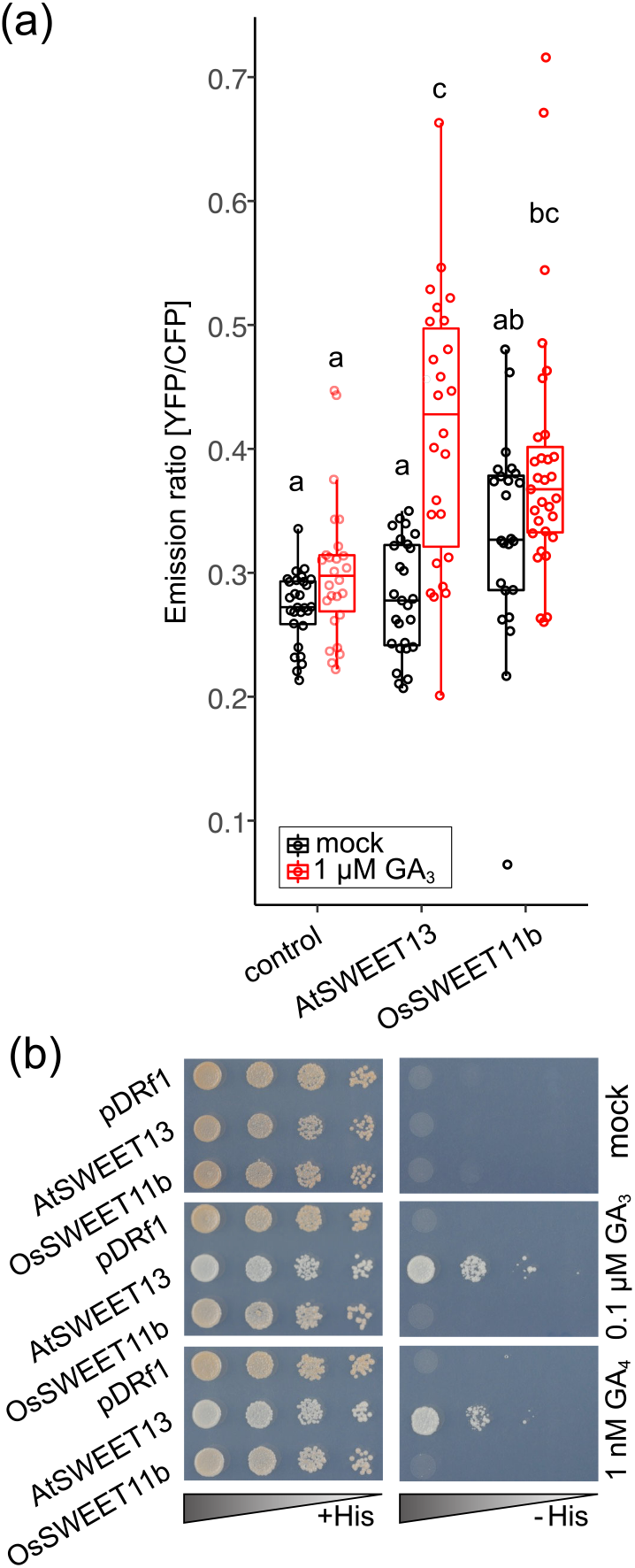
Gibberellin transport activity of OsSWEET11b. (a) 1 μM GA_3_ was added for 3 h to HEK293T cells expressing the GA sensor GPS1. Empty vector and AtSWEET13 served as negative and positive controls, respectively. Box plots show first and third quartiles, split by the median with whiskers extending 1.5x interquartile range beyond the box. Each data point represents mean fluorescence ratio during 3 min recordings (*n* = 28; empty vector, mock), 26 (empty vector, 1 μM GA_3_), 27 (*AtSWEET13*, mock), 26 (*AtSWEET13*, 1 μM GA_3_), 31 (*OsSWEET11b*, mock), 24 (*OsSWEET11b*, 1 μM GA_3_) cells. Two independent replicates were conducted (Fig. S16). Different letters on each boxplot represents significant difference determined by One-way ANOVA with Tukey’s post-hoc test (*p* < 0.05). (b) GA_3_ and GA_4_ transport activity of OsSWEET11b assessed using a GA-dependent Y3H system. Yeast strain PJ69-4a carrying both pDEST22-GID1a and pDEST32-GAI and either pDRf1-OsSWEET11b, pDRf1-AtSWEET13 (positive control), or empty vector (negative control) were grown on SD (-Leu, -Trp, -Ura) or selective SD(-Leu, -Trp, -Ura, -His) medium, containing 3 mM 3-AT and 0.001% (v/v) DMSO and either 0.1 μM GA_3_, or 1 nM GA_4_, respectively, for 3 days at 30°C. Comparable results were obtained in three independent replicates (Fig. S17).

### OsSWEET11b functions as a potential susceptibility gene for BLB

Up until now, naturally occurring TALes that target *OsSWEET11b* have not been described. An alignment of the promoter region showed that the binding site (EBE) for the TALe PthXo1 was only present in *OsSWEET11a*, indicating that *OsSWEET11b* is not targeted by PthXo1 from PXO99^A^ (Fig. S5b). The promoter of *OsSWEET11b* did not contain binding sites for other known TALes either. Designer TALes can be expressed in disarmed *Xoo* to test whether SWEET genes can function as susceptibility genes. Three of the five previously known clade III SWEET family members are currently targeted by the known *Xoo* strains from Asia and Africa, however all five can function as susceptibility factors when artificially induced by *Xoo* strains producing designer TALes (Streubel *et al*., 2013; Oliva *et al*., 2019). To test whether *OsSWEET11b* can also serve as a potential susceptibility gene for *Xoo*, a designer dTALe that can bind to the TATA box region of the *OsSWEET11b* promoter in the *japonica* rice cultivar Kitaake was synthesized (Fig. S18). The synthetic dTALe gene, which encodes twenty-three 34-amino acid repeats was introduced into a disarmed *Xoo* strain ME2, a PXO99^A^ mutant in which the TALe *pthXo1* was inactivated and which lacks other TALe for *SWEET* gene induction. Infection of Kitaake with ME2(dTALe) strongly induced *OsSWEET11b* as shown by RT-PCR (Fig. 8a, Fig. S19) and analysis of GUS activity in OsSWEET11b-Gus lines (#1 and #13) (Fig. 8b). Successful infection occurred with ME2(dTALe) in two *japonica* and *indica* rice cultivars Kitaake and Guanglu’ai 4, as dTALe provided ME2 full virulence (Fig. 8c, d). In summary, *OsSWEET11b* could be activated by dTALe and thus represents a potential BLB susceptibility that is likely related to sucrose transport activity of OsSWEET11b.

**Figure 8.**
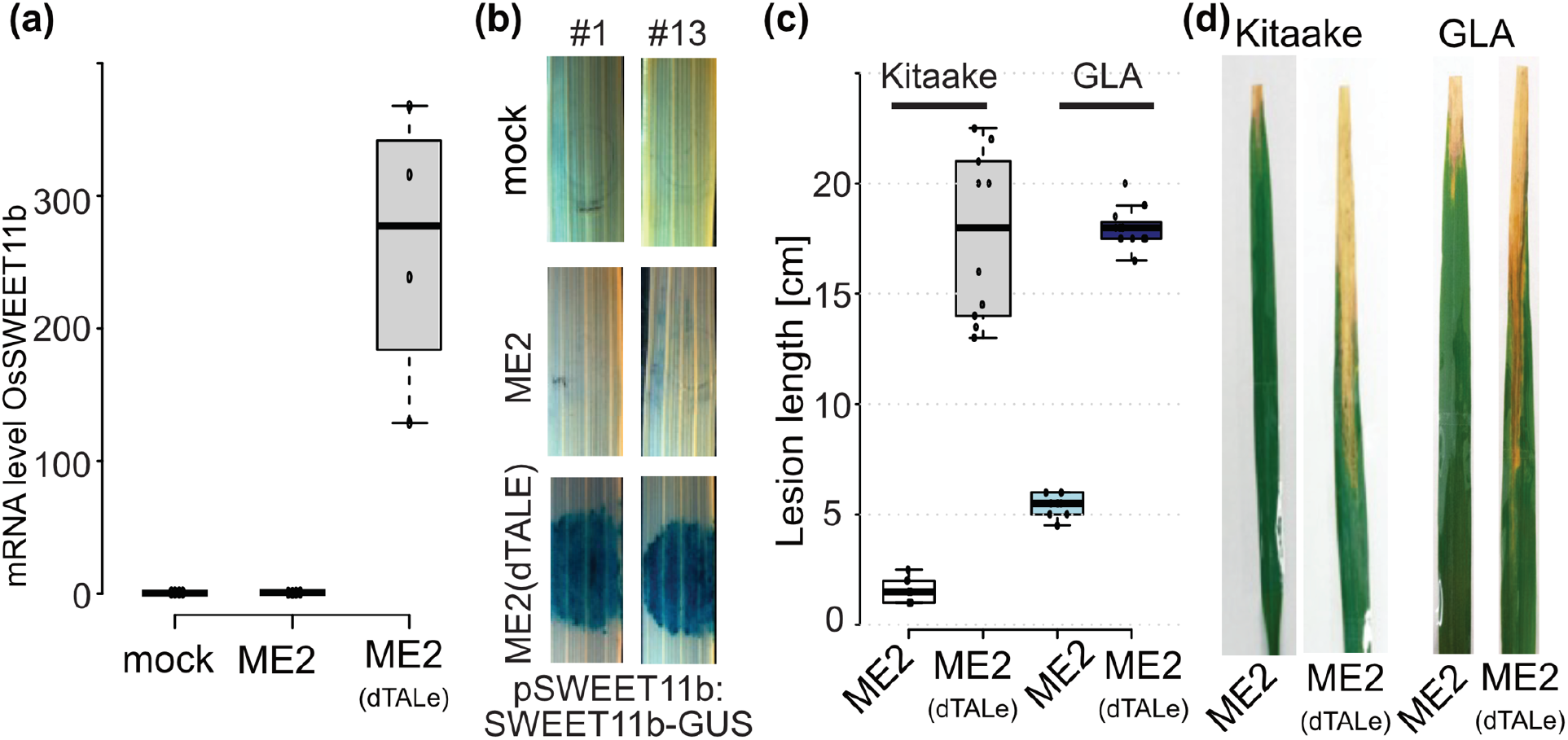
*OsSWEET11b* is a susceptibility gene for rice bacterial blight. (a) dTALe induces *OsSWEET11b*. mRNA level of *OsSWEET11b* (2^−ΔΔCt^) determined by qRT-PCR in Kitaake leaves treated with water (mock inoculation), ME2 and ME2 containing designer TALe. The rice actin gene was used as an internal control. (b) GUS histochemistry of rice leaves expressing translational OsSWEET11b-GUS fusions after inoculation with ME2 and ME2(dTALe). Results from two independent transgenic lines (#1; #13). (c) Designer TALe (dTALe) in ME2 causes virulence in rice. Lesion length measurements caused by *Xoo* strains ME2 and the designer TALe strain ME2/dTALe in two rice cultivars Kitaake (*japonica* rice variety) and Guanglu’ai 4 (GLA, *indica* rice variety) 14 days post inoculation. (d) Images of leaves showing lesions of blight with leaf tip clipping inoculation method. Experiments were repeated at least three times independently.

## Discussion

Here, a new member of the SWEET gene family was identified and named *OsSWEET11b*, based on its close phylogenic relation to OsSWEET11a (formerly named OsSWEET11 or Os8N3). This gene had not been detected in earlier genome annotations; interestingly in new annotations of the maize genome the close homolog ZmSWEET11b was also newly found. OsSWEET11b protein accumulated in anther veins, while OsSWEET11a was present in veins of stamina that enter the anthers (anther peduncle). Single mutants had no apparent sterility defects, however, when combined, double mutants became fully male sterility. The sterility is likely due to a defect in sucrose supply to the megaspores during their development, resulting in insufficient reserves for germination. While the Arabidopsis homologs AtSWEET13 and 14 may play roles in supplying GA to support fertility, OsSWEET11b showed sucrose transport activity, but no detectable GA transport activity, and GA did not supplement the infertility. The ability of SWEETs to recognize and transport substrates as diverse as sucrose and GA remains enigmatic. At present, it cannot be excluded that the observed GA transport activity is physiologically not relevant, and GA supplementation of sterility in Arabidopsis is due to indirect effects. Surprisingly, the evolution of GA transport capacity by SWEETs appears to emerge or gets lost during evolution independently (Morii *et al*., 2020). Dissection of the two activities by combination of structural and mutagenesis studies and further characterization of SWEETs may help to resolve this conundrum.

The use of designer TALe in disarmed *Xoo* strains shows that *OsSWEET11b*, similar to its paralog *OsSWEET11a*, can serve as a susceptibility locus. However, at the current point of time, there are no known *Xoo* strains that target *OsSWEET11b* to cause BLB. Originally, hairpin RNAi with a 598 bp fragment corresponding to the 3’-end of *Os8N3* (renamed *OsSWEET11a* herein) caused reduced male fertility (Yang *et al*., 2006). Notably, the starch content of pollen grains was reduced as seeb here. However, surprisingly, *ossweet11a knock out* lines did not show obvious male fertility defects in greenhouse conditions (Yang *et al*., 2018). The defects in the RNAi lines were thus likely due to RNA interference impacting both *OsSWEET11a* and *11b*.

### Interplay of SWEET11a and b with other sugar transporters and cell wall invertases

We initially hypothesized that OsSWEET11a and 11b would be responsible for the release of sucrose from the tapetum and uptake into the developing megaspores. Unexpectedly, both proteins were found in veins of the anther, with perfectly complementary localization in the peduncle of anther (OsSWEET11a) and the adjacent veins along the axis of the anther (OsSWEET11a). This clear separation may indicate that there are multiple apoplasmic steps, one at the interface between peduncle and anther, and another between the phloem and the connective. To our knowledge, there are at present no data regarding symplasmic domains in rice that are based on dye coupling studies or GFP movement as for *Arabidopsis*. However, while in *Arabidopsis* GFP can enter the anther tissues, there appears to be a step gradient, possibly implying a constriction that limits GFP entry leading to much higher fluorescence at the anther peduncle of Arabidopsis (Imlau *et al*., 1999). Dye coupling and GFP movement studies may help to unravel the exact pathways. SWEETs are uniporters that can either enable efflux of sucrose or function in cellular uptake, dependent on the sucrose concentration gradients. Apoplasmic sucrose released from cells can either be taken up into adjacent cells by SWEET - or SUT-mediated uniport or hydrolyzed by cell wall invertases followed by uptake via MST hexose transporters (Goetz *et al*., 2017). Sucrose released by the SWEETs may subsequently in part be taken up by a SUT sucrose transporter; in Arabidopsis AtSUC1 produced in a cell layer of the connective adjacent to the vasculature is a candidate (Stadler *et al*., 1999). Genes for cell wall invertases INV1 and INV4 are both expressed in maturing rice anthers, however, INV4 is transiently present in the tapetum and later in microspores and may play a role in cold sensitivity (Oliver *et al*., 2005). Consistent with extracellular hydrolysis, two hexose/H^+^ symporters *MST7* and *8* seem to be coexpressed with the invertases. These findings implied the presence of sucrose efflux transporters that supply the invertase with sucrose. Clade III SWEETs are prime candidates for such functions. The presence of cell wall invertases in multiple locations in the anther, tapetum and developing microspores matches the patterns of SWEET11a and b accumulation.

Several regulatory proteins have been identified that are involved in regulation of MSTs, invertases and several other transporters of the SUT and TMT family, indicating parallel pathways for apoplasmic flux of hexoses and sucrose. Notably, the MYB domain protein Carbon Starved Anther CSA, required for carbon supply and male fertility in rice, shows similar expression patterns as *OsSWEET11b. csa* mutants were characterized by reduced *MST8*, *SUT3* and *INV4* mRNA levels (Zhang *et al*., 2010). The bZIP transcription factor 73, which is required for cold tolerance in anthers, also plays a role in regulating *MST7*, *8* and *INV4* genes (Liu *et al*., 2019). Several sugar transporters have been shown to be regulated posttranscriptionally by the RNA binding protein OsRRM. OsRRM is produced in anthers and has been shown to positively affect steady state transcript levels of a suite of transporter genes including *MST6* and *8*, *TMT1* and *2*, as well as four sucrose transporters *SUT1*,*2*, *3* and *4*, likely by stabilizing the respective mRNAs (Liu *et al*., 2020). SWEET mRNA and protein levels have not been tested in these mutants, yet based on the findings summarized above, it is tempting to speculate that *SWEET11a* and *b* may be coregulated by the same mechanisms and may also be involved in temperature tolerance of male reproductive organs in rice.

While OsSWEET11b is a close homolog of OsSWEET11a, it has diverged sufficiently to not contain an effector binding site for PthXo1. Thus, *OsSWEET11b* cannot be targeted by PXO99^A^. A systematic analysis of the ability of SWEET gene family members to serve as BLB susceptibility factors using designer TALes had shown that induction of any of the five previously known clade III SWEETs supported virulence (Streubel *et al*., 2013). It is conceivable that *OsSWEET11b* cannot serve as a susceptibility gene because it is guarded (Jones & Dangl, 2006). However, here we show that *OsSWEET11b* can serve as a susceptibility gene to *Xoo* as long as a strain lacking suitable TALe for SWEETs is equipped with a designer TALe that bind to the *OsSWEET11b* promoter. Thus, while all six clade III SWEETs are potential susceptibility genes, only three out are used by the extant Asian and African *Xoo* strains. The finding that only half of the now six clade III SWEET susceptibility factors are actually used by the known *Xoo* strains indicates a limited ability of *Xoo* to evolve new TALes that target these promoters. This observation is in line with the strict continental isolation, where *Xanthomonas oryzae* strains from the Americas do not target any of the SWEETs and are weak pathogens causing limited damage (Triplett *et al*., 2011). African strains, target two regions in a single SWEET, namely *OsSWEET14*. Asian strains appear more ‘advanced’ in that they target three SWEETs (*OsSWEET11a*, *13* and *14*) with a diversified TALe portfolio and a series of adaptations to the allelic promoters in different rice varieties (Oliva *et al*., 2019). This ‘advanced’ state is likely due to Asia being by far the largest cultivation area for rice, and the origin of resistance breeding. Based on the analysis of *Xoo* strain collections from Asia and Africa it has been possible to develop elite rice varieties resistant to all strains in a representative collection (Oliva *et al*., 2019). Since it is conceivable that new strains evolve to target the still unused clade III SWEETs including *OsSWEET11b*, diagnostic tools will help to accelerate the discovery of emergent disease mechanisms and enable rapid discovery of SWEET promoter sequences targeted by TALes (Eom *et al*., 2019; Li *et al*., 2019). Based on the findings here it planned to expand the diagnostic kit and to accelerate the genome editing based generation of promoter variants for emerging EBEs in *OsSWEET11b*.

## Acknowledgements

We thank Bi Huei Hou for excellent technical assistance and sensor-based transport studies, and Bo Liu for rice transformation of CRISPR and GUS-reporter lines. We thank Mitsunori Seo (RIKEN, Yokohama, Japan) for advice and plasmids for the Y3H system components. We thank Michael Frei (Justus-Liebig University Giessen, Germany) for the support on experiments conducted at University Giessen. This work was supported by grants from the Bill & Melinda Gates Foundation to HHU (OPP1155704; WF), with a subcontract to MU (BY), the Alexander von Humboldt Professorship (WF), Deutsche Forschungsgemeinschaft (DFG, German Research Foundation) under Germany’s Excellence Strategy – EXC-2048/1 – project ID 390686111, a Japan Society for the Promotion of Science (JSPS) grant–19H000932 (WF, MN) and the World Premier Institute Center Initiative, Japan. Work in BYs lab was also supported by the National Science Foundation (IOS-1936492).

## Author contributions

WBF, BY and MN conceived of the study, designed experiments, and supervised the work. WBF, BY and LBW wrote the manuscript. JSE generated GUS constructs. VTL performed GFP fusion localization assays, LBW performed GUS assays, genotyping and phenotyping plants and in planta GA assays, LC carried out gene induction and disease assays, SNC designed and constructed dTALe, generated DL mutants and carried out fertility analyses; RI performed all GA transport assays.

## Data Availability

The data that support the findings of this study are available from the corresponding author upon reasonable request.

