## Supplementary Data for "*OsSWEET11b*, a sixth leaf blight susceptibility gene involved in sugar transport-dependent male fertility"

**Contents:**

Fig. S1 Construct of CRISPR-Cas9 vector targeting *OsSWEET11b*
Fig. S2 Construct of CRISPR-cas9 vector simultaneously targeting *OsSWEET11a* and *11b*
Fig. S3 Construct for the translational fusion of OsSWEET11b with GUS under control of its own promoter
Fig. S4 Genotypic variations of two independent lines of *ossweet11b* single and *ossweet11a*;b double mutant.
Fig. S5 Rice clade III SWEET protein sequences analyses and OsSWEET11b transmembrane domain prediction
Fig. S6 Gene features of *OsSWEET11a* and *11b*
Fig. S7 Transcript level of rice clade III SWEETs in various organs at the reproductive stage
Fig. S8 OsSWEET11b-GUS histochemistry in rice seedlings and spikelet branches
Fig. S9 Localization of OsSWEET11b protein in *N. benthamiana* leaves
Fig. S10 Subcellular localization of OsSWEET11b
Fig. S11 Sucrose transport by OsSWEET11b
Fig. S12 Phenotype of panicles and grains in *ossweet11a, 11b single and ossweet11a;b double mutant* in Düsseldorf greenhouses
Fig. S13 Carpels of wild type Kitaake, *ossweet11a* and *ossweet11b* mutants
Fig. S14 $F_1$ seeds from reciprocal crosses between *ossweet11a;b-1*, *ossweet11a-2* (a, b) and *ossweet11a;b-1*, *ossweet11b-1*
Fig. S15 Florets of wild type Kitaake, *ossweet11a-1*, *ossweet11b-2* and *ossweet11a;11b-2* mutant lines
Fig. S16 $GA_3$ uptake by OsSWEET11b into human Hek293T cells
Fig. S17 Independent experiment showing $GA_3$ or $GA_4$ uptake by OsSWEET11b into yeast cells
Fig. S18 Gene structure of Os*SWEET11b* with sequences shown for designer TALe and CRISPR/Cas9 guide RNA
Fig. S19 Induction of *OsSWEET11b* mRNA level in rice infected with different Xoo strains
Table S1 Primers used in this study
Table S2 Genotype and phenotype analyses of $T_3$ plants of *ossweet11a,b* double mutants

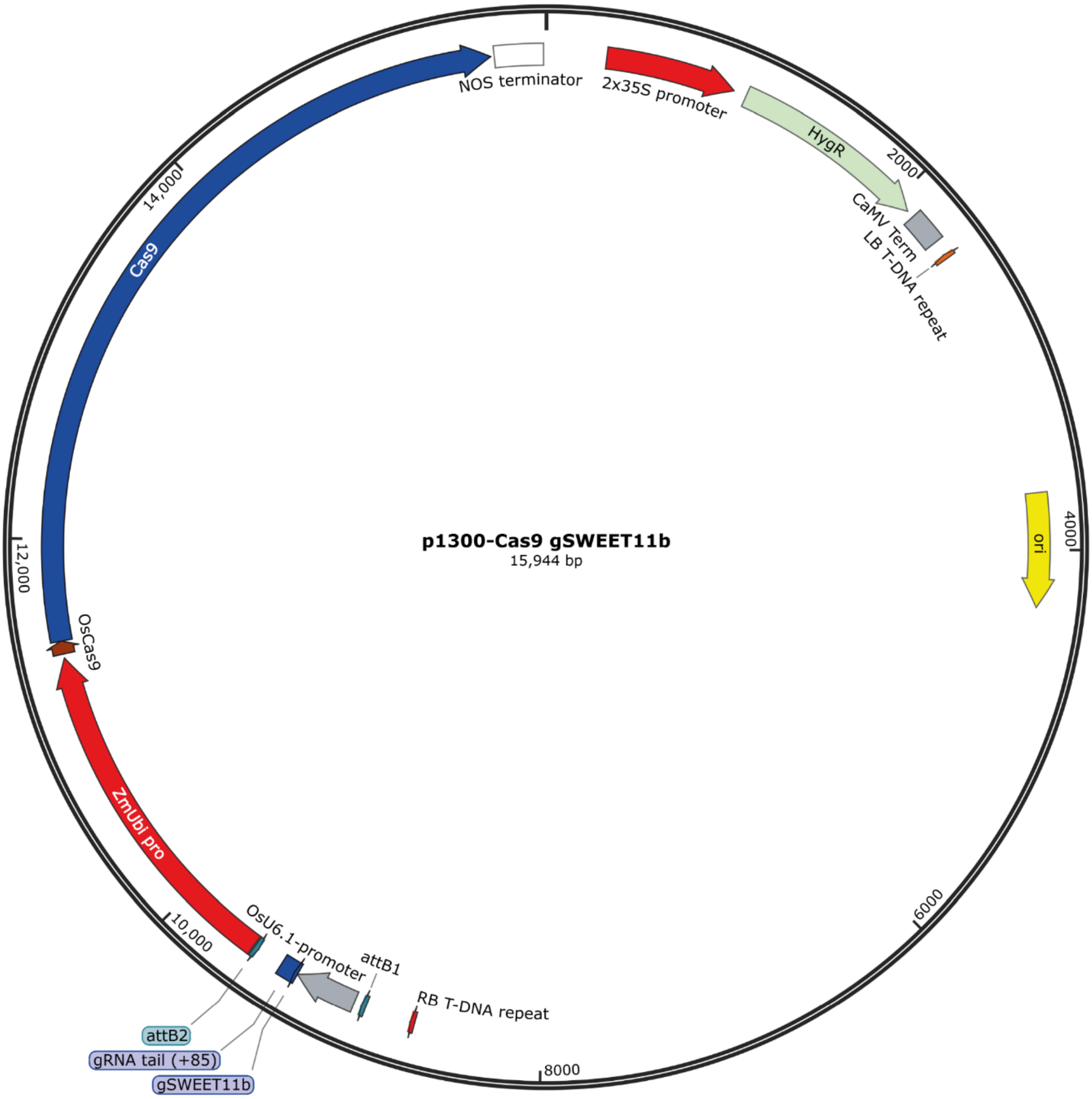


**Figure S1. Construct of CRISPR-Cas9 vector targeting *OsSWEET11b*.** Transcription of the *OsWEET11b*-trageting guide RNA is driven by the OsU6.1 promoter. Cas9 gene was driven by the *ZmUbi* promoter.

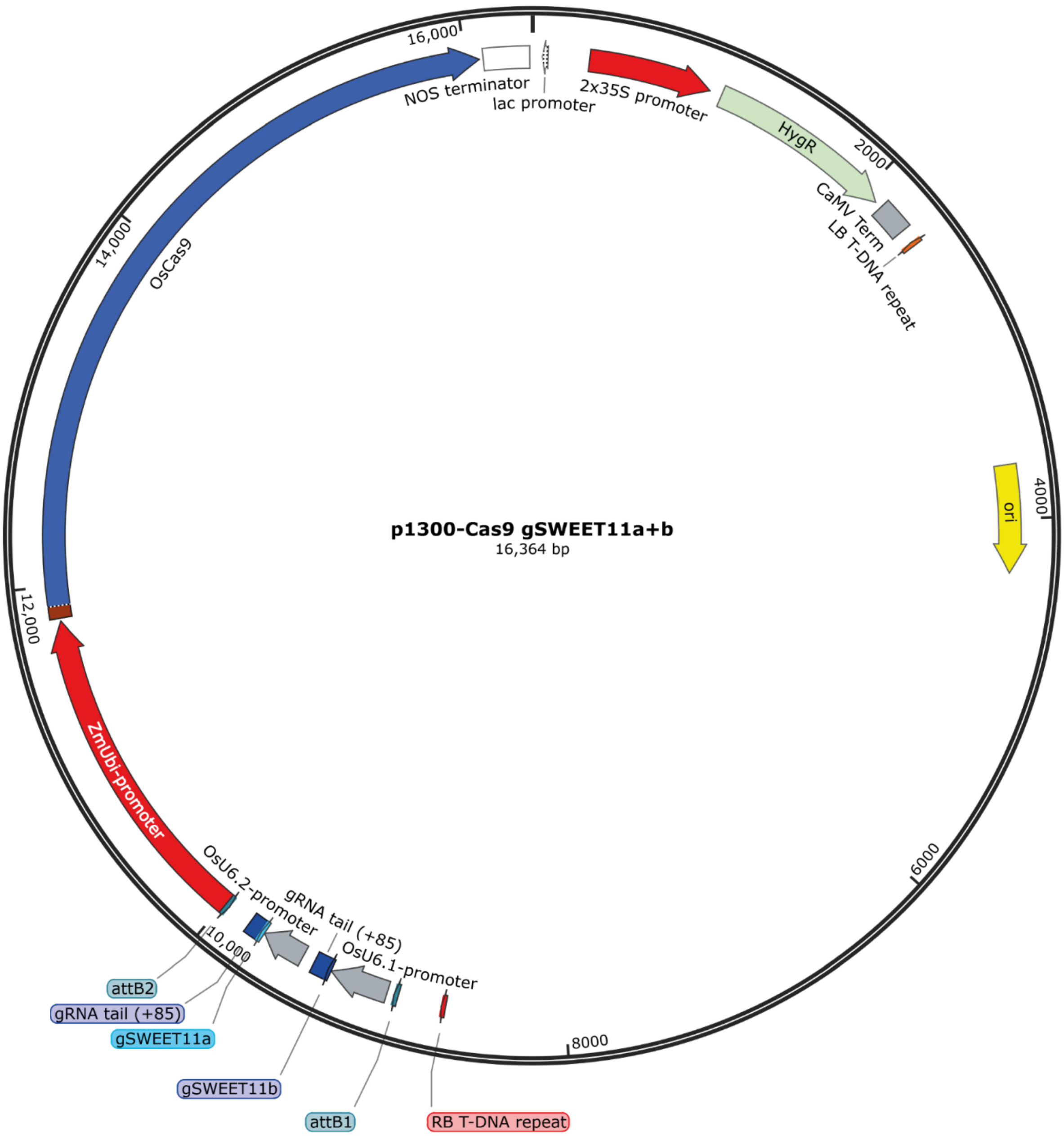


**Figure S2. Construct of CRISPR-cas9 vector simultaneously targeting *OsSWEET11a* and *11b*.** Transcription of *OsWEET11b* and *11a* single guide RNA driven by OsU6.1 and OsU6.2 promoter, respectively. Cas9 gene was driven by *ZmUbi* promoter.

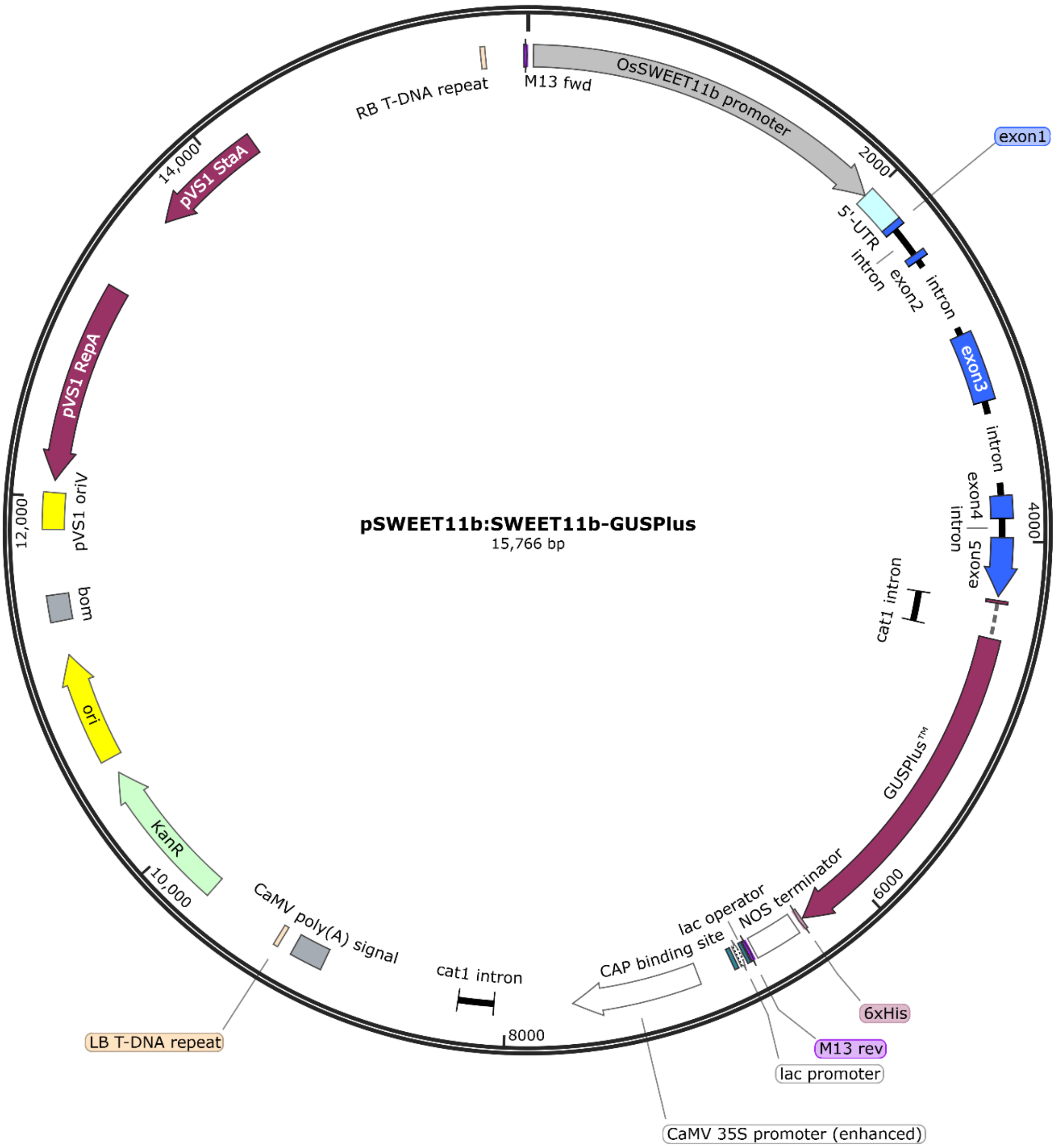


**Figure S3. Construct for the translational fusion of OsSWEET11b with GUS under control of its own promoter.** A genomic fragment of 4,262 bp in *OsSWEET11b* including 2155 bp of promoter region and all introns and exons (without stop codon) was introduced into promoterless GUSPlus vector.

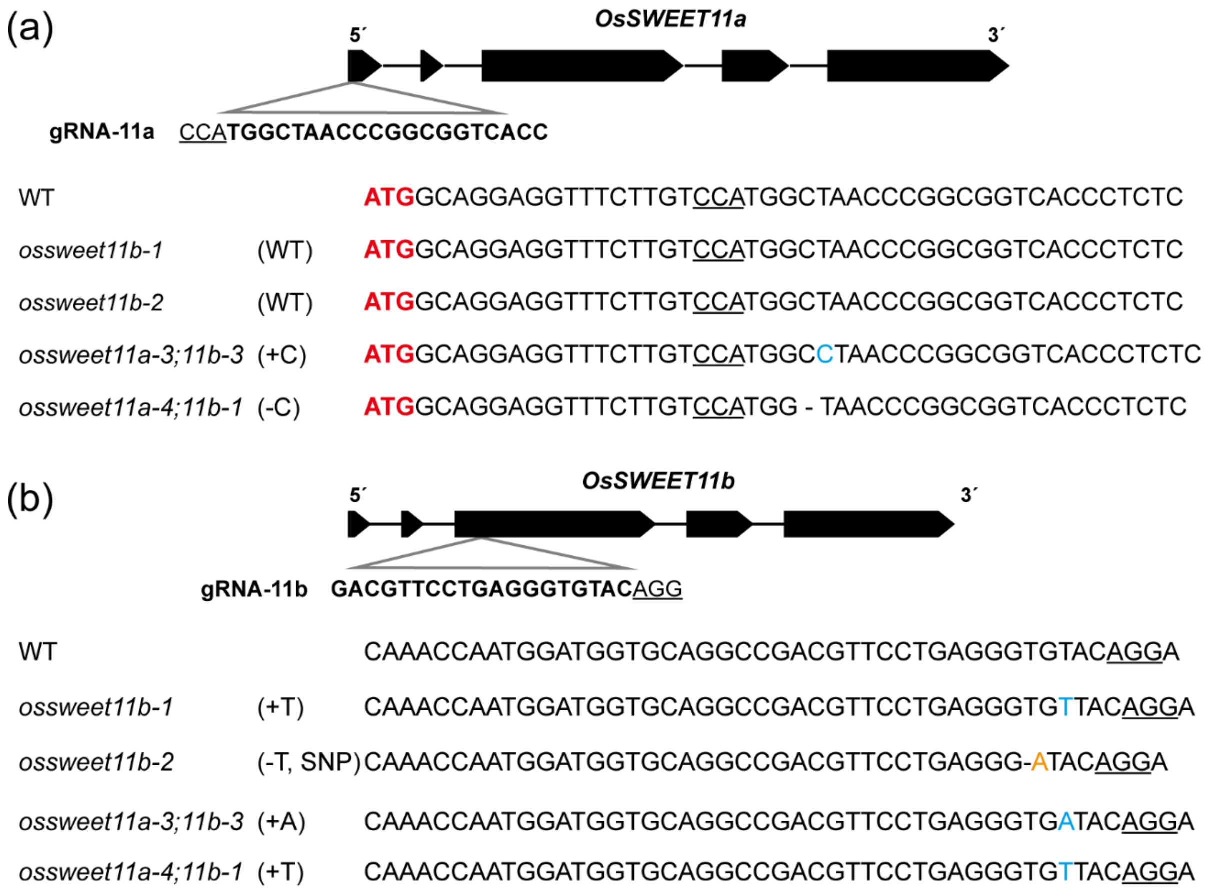


**Figure S4. Genotypic variations of two independent *ossweet11b* single and *ossweet11a*;b double mutants.** (a) Single guide RNA and PAM sequences (underlined) targeting the first exon in *OsSWEET11a* resulting different types of mutations in *ossweet11a;b* double mutant lines. Start codons were marked red and single nucleotide insertion was marked blue. (b) Single guide RNA and PAM sequences (underlined) targeting the third exon in *OsSWEET11b*. Single nucleotide insertions were marked blue and SNP was marked orange.

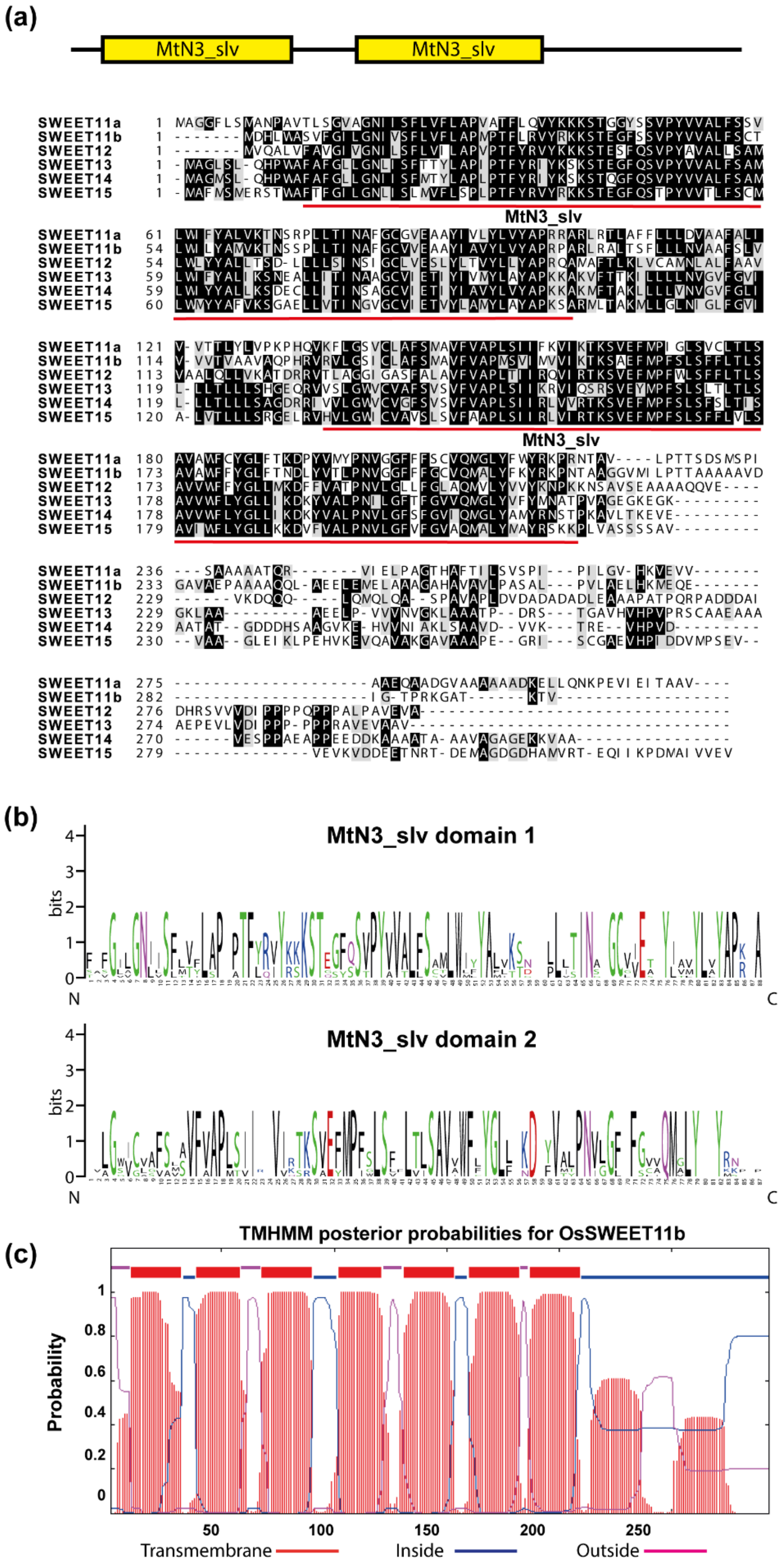


**Figure S5. Rice clade III SWEET protein sequences analyses and OsSWEET11b transmembrane domain prediction.** (a) Alignment of all six clade III SWEET protein sequences. **(b)** Sequence logo of clade III SWEET proteins (https://weblogo.berkeley.edu). **(c)** Transmembrane domain prediction using TMHMM V2 (http://www.cbs.dtu.dk/services/TMHMM/).

(a)

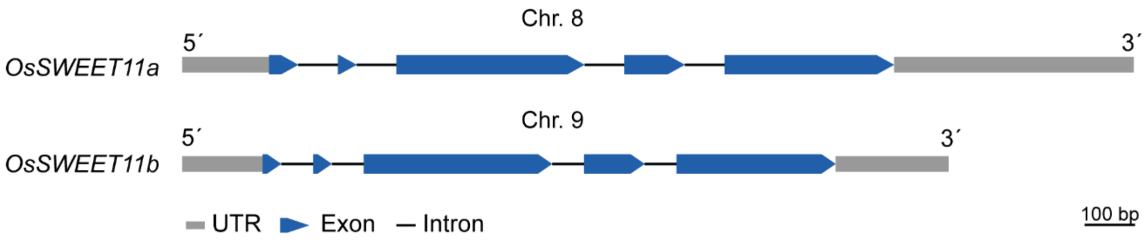


(b)

```
OsSWEET11b_pro      CTCTACAATATCTAAATATGTCTCTCCTATGTACTAACTTTTTGAATACATTCCTGAATT
OsSWEET11a_pro      --------TATGACCA-----------CATATTCAGAGTAGT-GGAGAGAGGGACAGATC
                            ***    *            ** * *  * *  * * * * *       **

OsSWEET11b_pro      --TAGTTAATGGAAGGATTATTATCTGACAAATGTTCAATTATTCAGTAAGA----CTTC
OsSWEET11a_pro      TAGAGGTAGAAAAAGAAAATTCA---TATAAATGATATATCAGAGTGAAAAAGAAATATC
                       ** **    *** *   * *    * ***** *  ** *    * ** *      **

OsSWEET11b_pro      AGGTTA--TAGCAAACAGTAGGAGTATAATATCTCTGGACCACTTAGGCACTTAGGCACC
OsSWEET11a_pro      AAGCACAAGAAAAAAAAGCAAAGGTTAGATATGCATCTCCCCCTACTGTAC---------
                    * *      *  *** ** *   **   ****   *   ** **   * **

OsSWEET11b_pro      ATGTAGATACGTACATATGTAACATATGAGAAGAAGAGAAGCTCTACTACTATATATACA
OsSWEET11a_pro      ----------------------ACCAC--CAAAAGTGGAGGGTCTCCAACTATATAAACA
                                             *    ** *   ** * *** * ******** ***

OsSWEET11b_pro      AGGCTTCTACCCCCAATGCCAAGCACACAAACATTGGGTGCAGTAGTGAATCCTC-----
OsSWEET11a_pro      CTGAGCCATGGCCAAGGCCAAACCACACATGCAGTTGTAGTAGCACTTAAGCCTTCCTCT
                      *   *    ** *   * ** ******  ** * *  * ** * * ** ***

OsSWEET11b_pro      -------------------TTCAGAAGTTCAGA--AAACTATAGCTTGCTTTTAGCATAT
OsSWEET11a_pro      CTAGCTAGCATCTCTTGTGTCAGGAAGTTGGAAGGGATTTCTGGCTAGTTTCTAGCTGGT
                                       *   ******   *   *  * * *** * ** ****   *

OsSWEET11b_pro      ATA-----------------AATTAGGTGTTTTTATCTTCTT-----TTGAGAAGGAGAT
OsSWEET11a_pro      GTCTCCTCTCCTCTTCCTAACCTTCTCACTGATTAACACCTTAGAGTTAGTTAATAACCT
                     *                    **     *  *** *  ***     * *  **  *  *

OsSWEET11b_pro      CAAGAACAGGCAGTAGGAGGCCTCTTCCTCTTCTCCATGGATCACCTTTGGGCCTCTGTC
OsSWEET11a_pro      TCATCACCAGTAGCAATGGCAGGAGGTTTCTTGTCCATGGCTAACCCGGCGGTCACCCTC
                      *  **  * ** *   *         **** ******* * ***    ** * *  **

OsSWEET11b_pro      TTTGGGATTCTT-
OsSWEET11a_pro      TCCGGTGTTGCAG
                    *  **  **
```

**Figure S6. Gene features of *OsSWEET11a* and *11b*.** (a) Conserved exon intron structure of *OsSWEET11a* and *11b*. (b) Alignment of 400 bp of promoter sequences of *OsSWEET11a* and *11b*. Start codons of *OsSWEET11a* and *11b* are highlighted with a red box, TATA-boxes in green and effector binding element (EBE) in *OsSWEET11a* which is targeted by the TALe PthXo1 from the *Xanthomonas oryzae* pv. *oryzae* strain PXO99[A] in yellow. The promoter sequences are poorly conserved and *OsSWEET11b* contains no sequences that can function as an EBE for PthXo1.

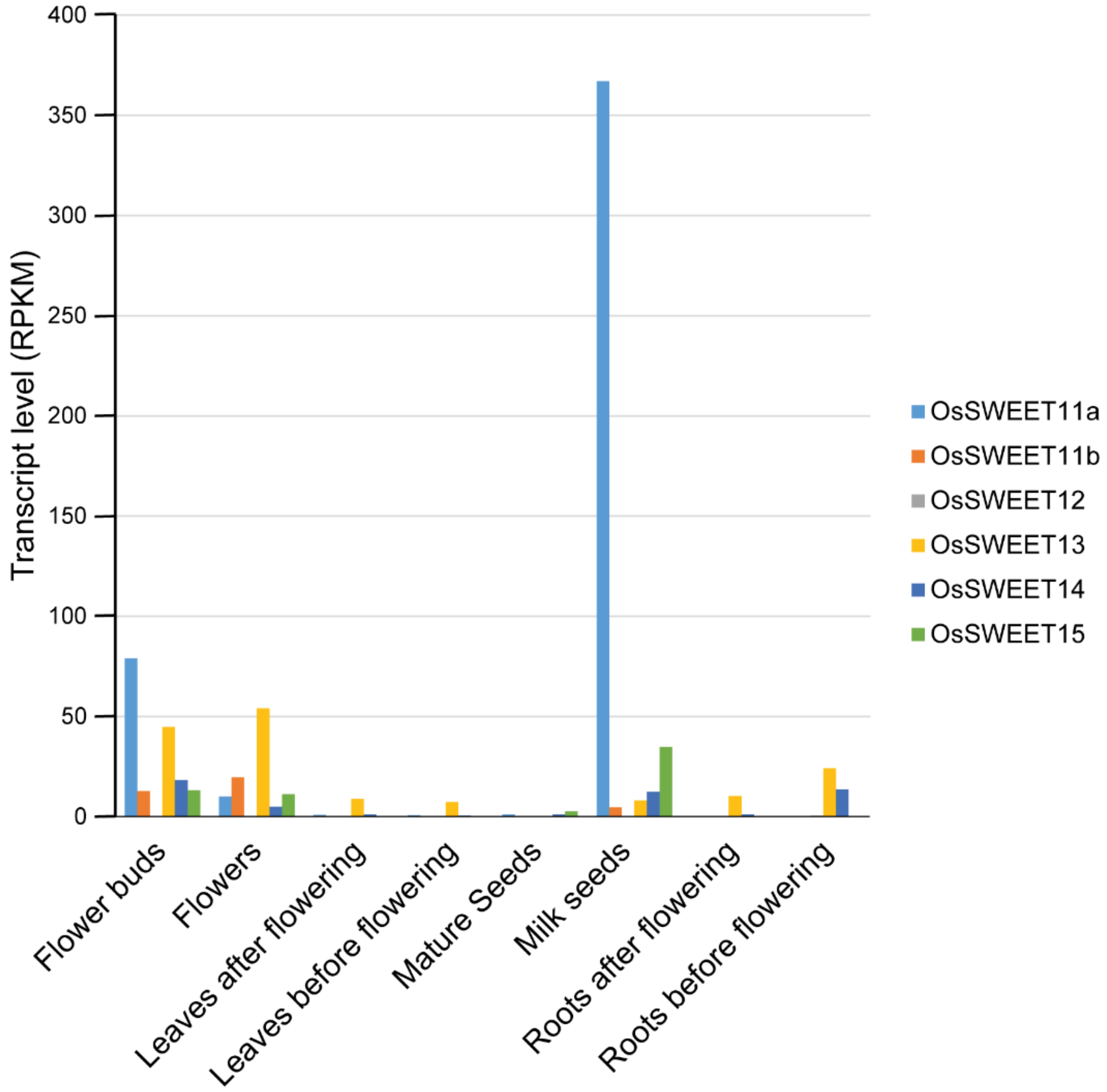


**Figure S7. Transcript level of rice clade III SWEETs in various organs at the reproductive stage.** RPKM values from Bioproject PRJNA243371 (https://www.ncbi.nlm.nih.gov/bioproject/PRJNA243371/) were collected from NCBI (Wang, Niu et al. 2015).

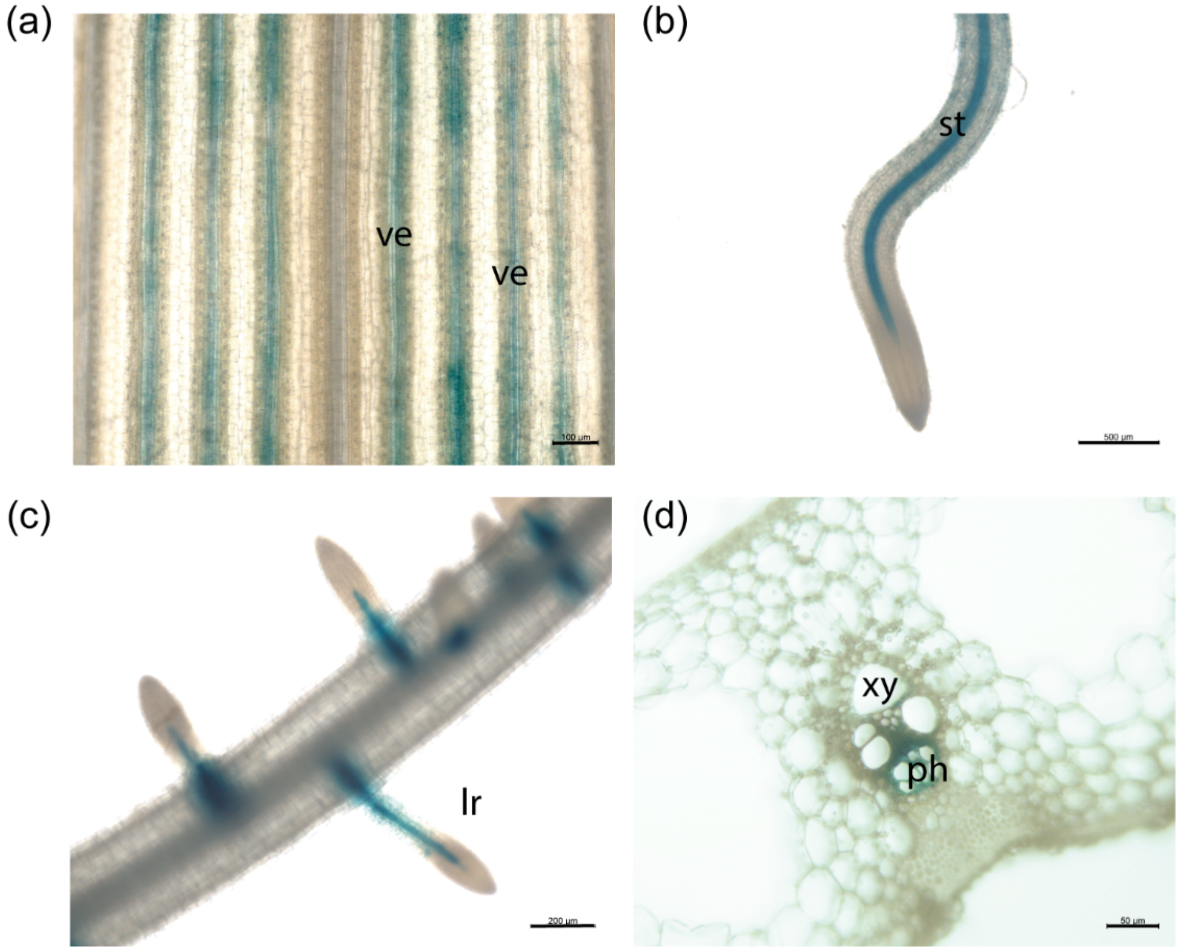


**Figure S8. OsSWEET11b-GUS histochemistry in rice seedlings and spikelet branches.**

OsSWEET11b-GUS activity was detected in leaf veins (ve), stele of primary root (st), lateral roots (lr) in 5-d-old rice seedlings and phloem (ph) of spikelet branches. Xy: xylem. Scale bars: (a) 100 µm; (b) 500 µm; (c), 200 µm and (d) 50 µm.

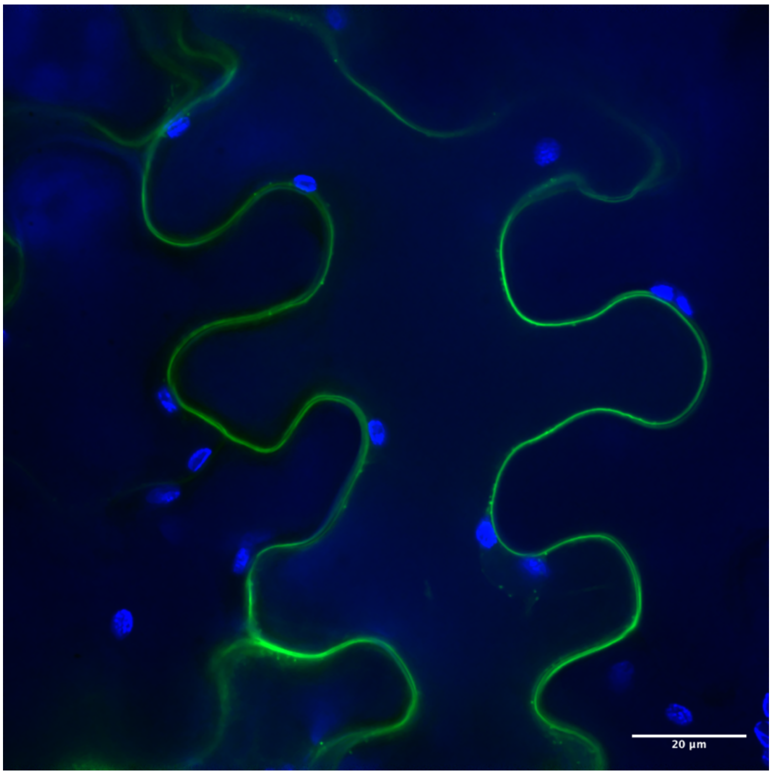


**Figure S9. Localization of OsSWEET11b protein in *N. benthamiana* leaves.** Confocal image of epidermal cells. A C-terminal fusion of eGFP to OsSWEET11b likely localizes to the plasma membrane (eGFP: green; chloroplast fluorescence: blue; 667–773 nm: blue). EGFP fluorescence was observed external to the peripheral chloroplasts excluding cytosolic and tonoplast localization. The experiment was repeated three times independently.

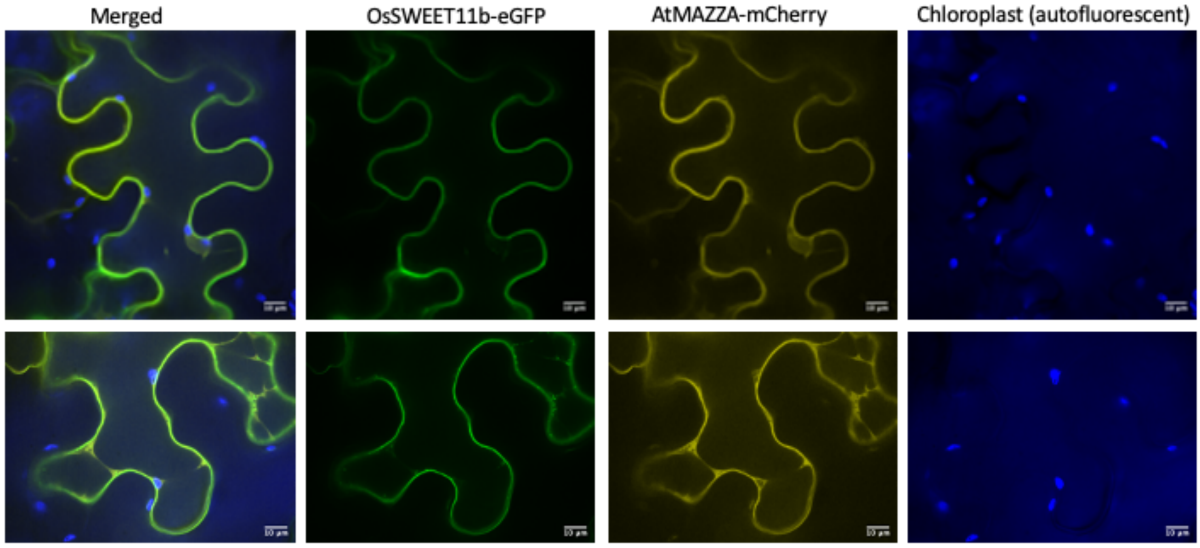


**Figure S10. Subcellular localization of OsSWEET11b.** A translational eGFP-fusion of OsSWEET11b was transiently expressed in *N. benthamiana* leaves and the subcellular localization was analyzed by confocal microcopy. Autofluorescence from peripheral chloroplasts in epidermal cells and the plasma membrane marker AtMAZZA-mCherry served as controls.

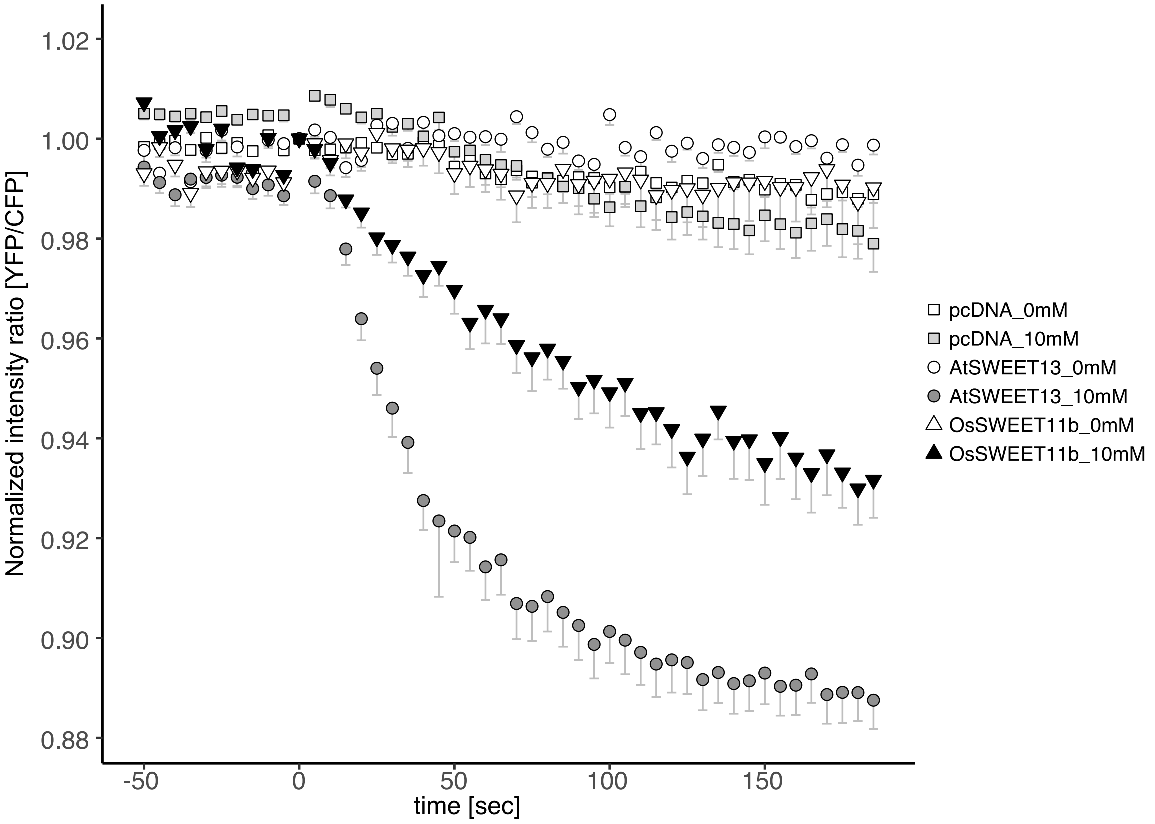


**Figure S11. Sucrose transport by OsSWEET11b.** Independent experiment for determining sucrose transport activity of OsSWEET11b in HEK293T cells using the sucrose sensor FLIPsuc90µ. HEK293T cells were co-transfected with constructs carrying FLIPsuc90µ and OsSWEET11b by Lipofectamine LTX (Invitrogen) in 8-well grass bottom chamber (Iwaki Cat#; 5232-008) and incubated for 48 h. Culture medium was replaced with 150 µL of HBSS buffer before the observation. The fluorescence was acquired under the same conditions as the GA uptake assays. 150 µL of HBSS buffer containing 20 mM sucrose was added during the observation. Empty vector (pcDNA3.1) and *AtSWEET13* (pcDNA3.2-AtSWEET13) served as negative and positive controls, respectively. The time of treatment was set to zero (mean - S.E.M, n ≥ 10).

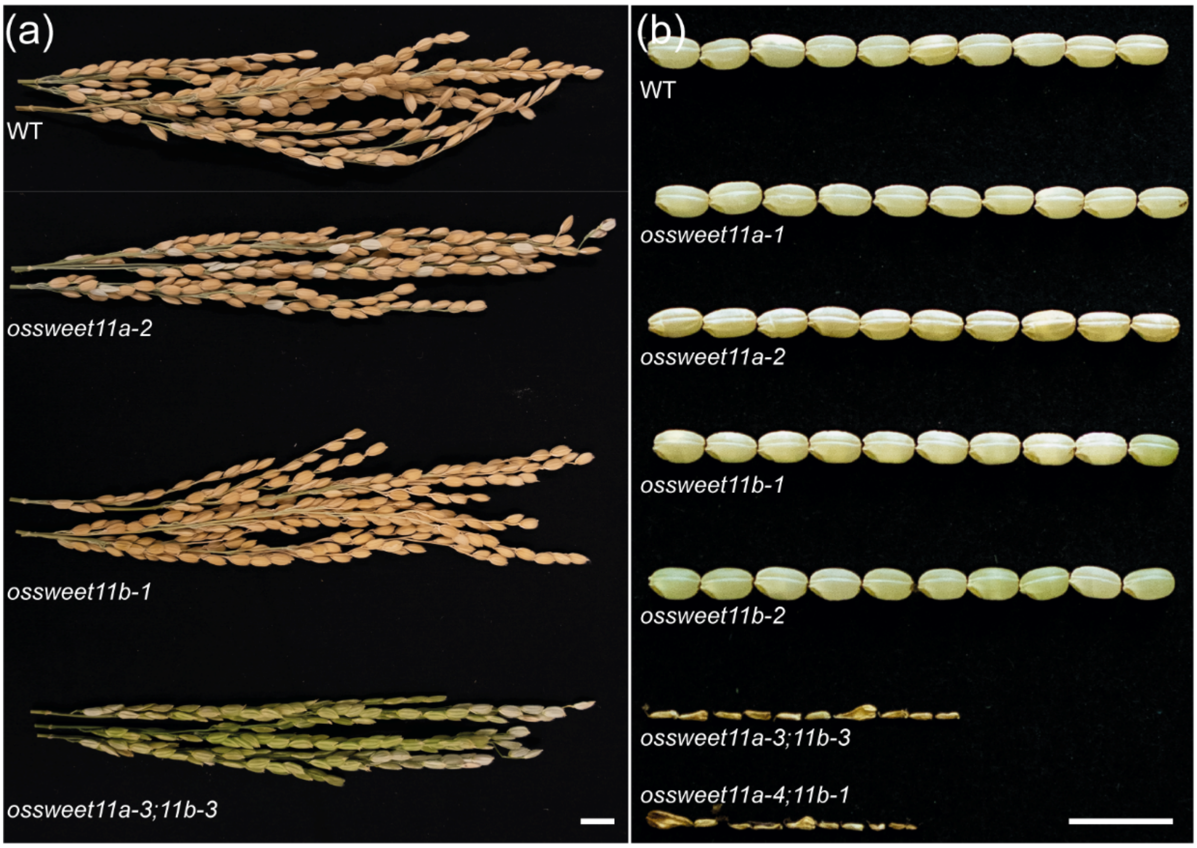


**Figure S12. Phenotype of panicles and grains in *ossweet11a, 11b single and ossweet11a;b double mutant* in Düsseldorf greenhouses.**

(a) Mature panicles of wild type Kitaake, *ossweet11a-2*, *ossweet11b-1* and *ossweet11a-3;b-3*. Neither *stay green* phenotypes nor obvious differences in seed filling were observed in *ossweet11a* mutants grown in Düsseldorf greenhouses. (b) Mature grains of wild type Kitaake, *ossweet11a* single mutant, *ossweet11b* single mutant and *ossweet11a-3;11b-3, ossweet11a-4;11b-1* double mutant plants. No differences in seed filling were observed in Iowa (Yang *et al.*, 2018). Seed filling defects were observed in greenhouses in Giessen (this work; Fig. 4), and had previously been observed in greenhouses in Cologne (Yang *et al.*, 2018), as well as in field experiments (Ma *et al.*, 2017).

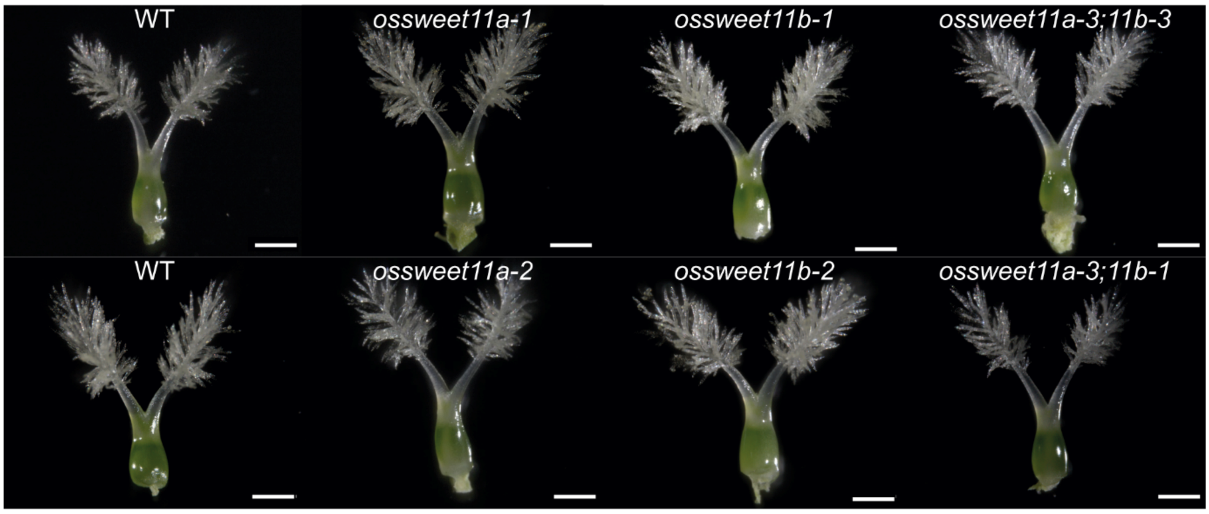


**Figure S13. Carpels of wild type Kitaake, *ossweet11a* and *ossweet11b* mutants.** Carpels show no obvious phenotypic differences between wild type Kitaake and *ossweet11a*, *ossweet11b* or *ossweet11a; 11b* double mutants. Scale bars: 200 µm.

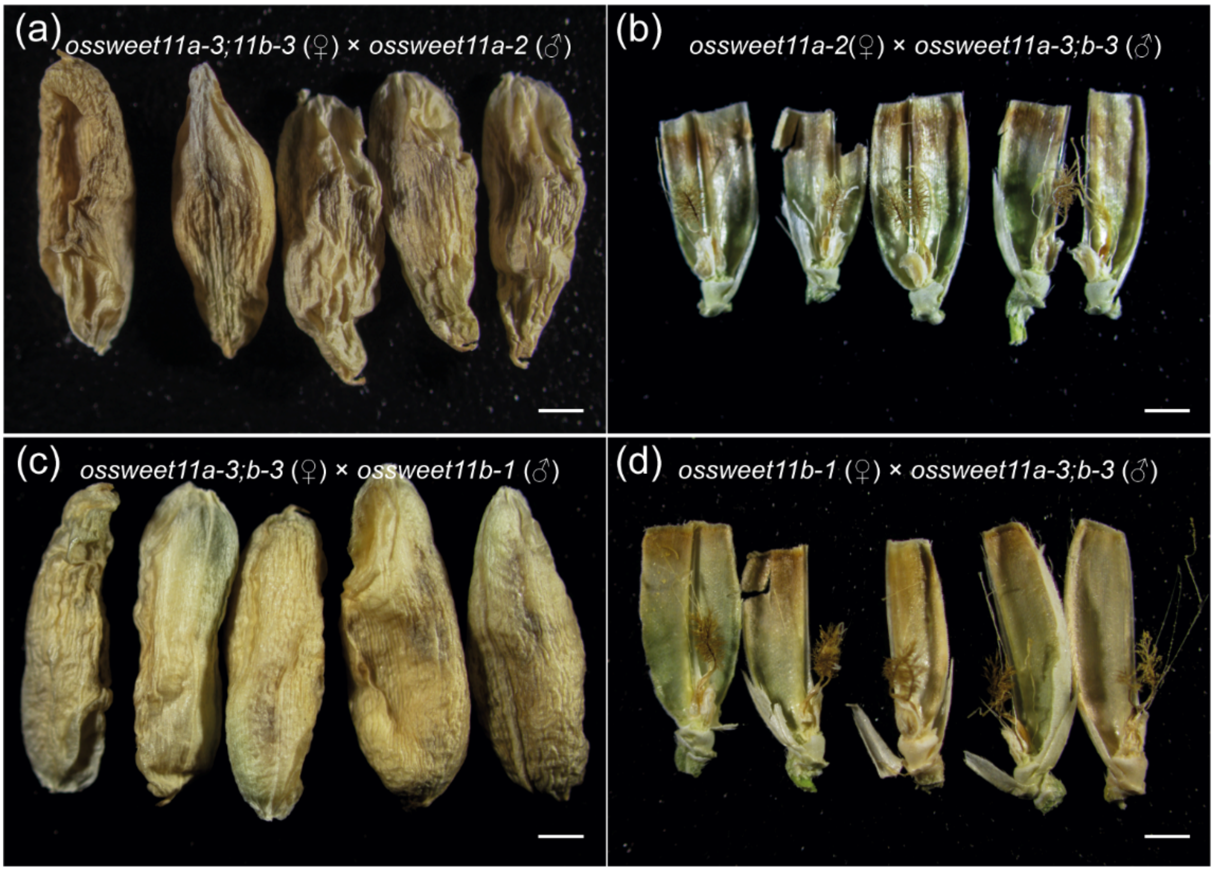


**Figure S14. $F_1$ seeds from reciprocal crosses between *ossweet11a-3;b-3*, *ossweet11a-2* (a, b) and *ossweet11a-3;b-3*, *ossweet11b-1* (c, d).** Scale bars: 1 mm.

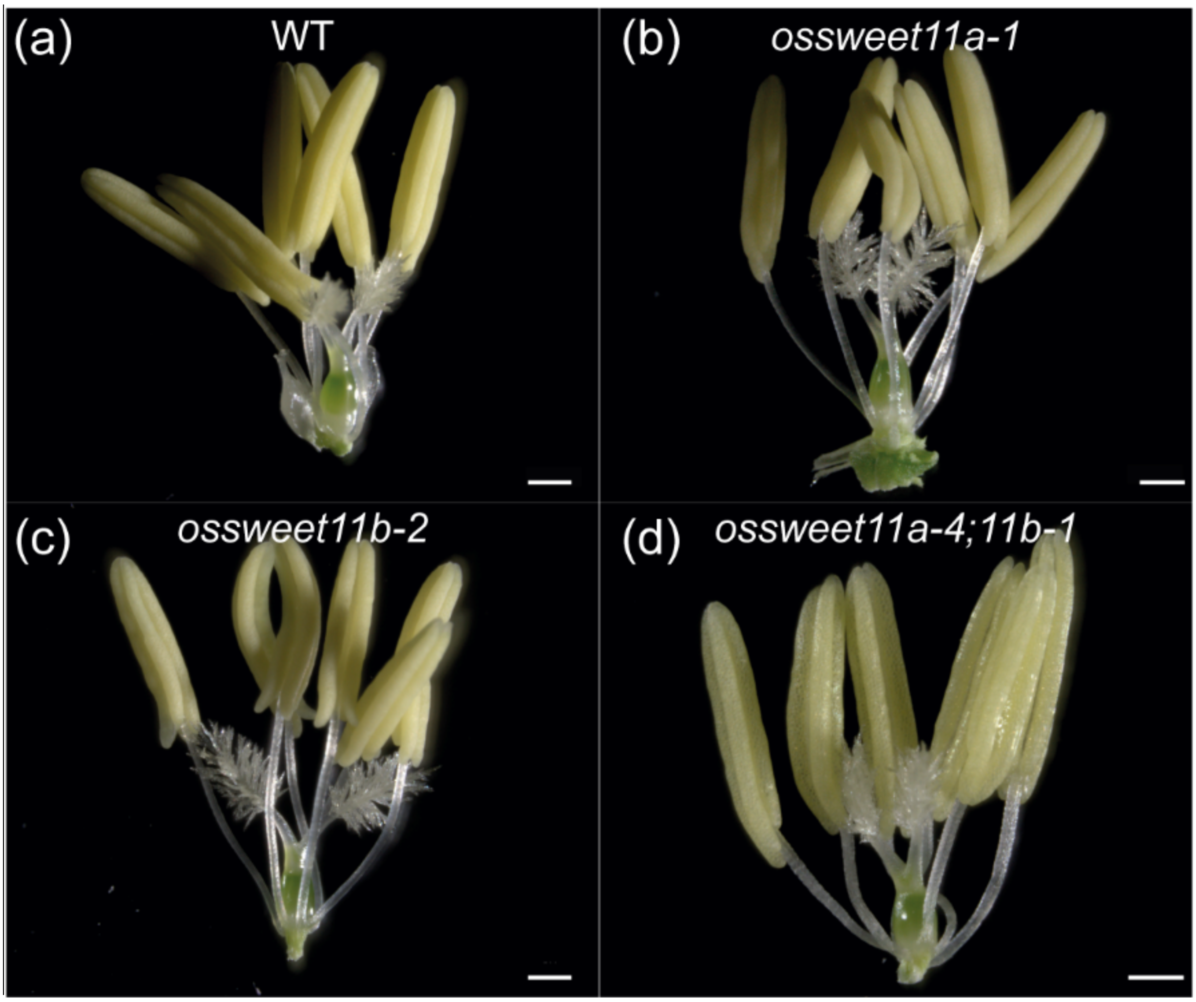


**Figure S15. Florets of wild type Kitaake (a), *ossweet11a-1* (b), *ossweet11b-2* (c) and *ossweet11a-4;11b-1* (d) mutant lines.** Scale bars: 500 µm.

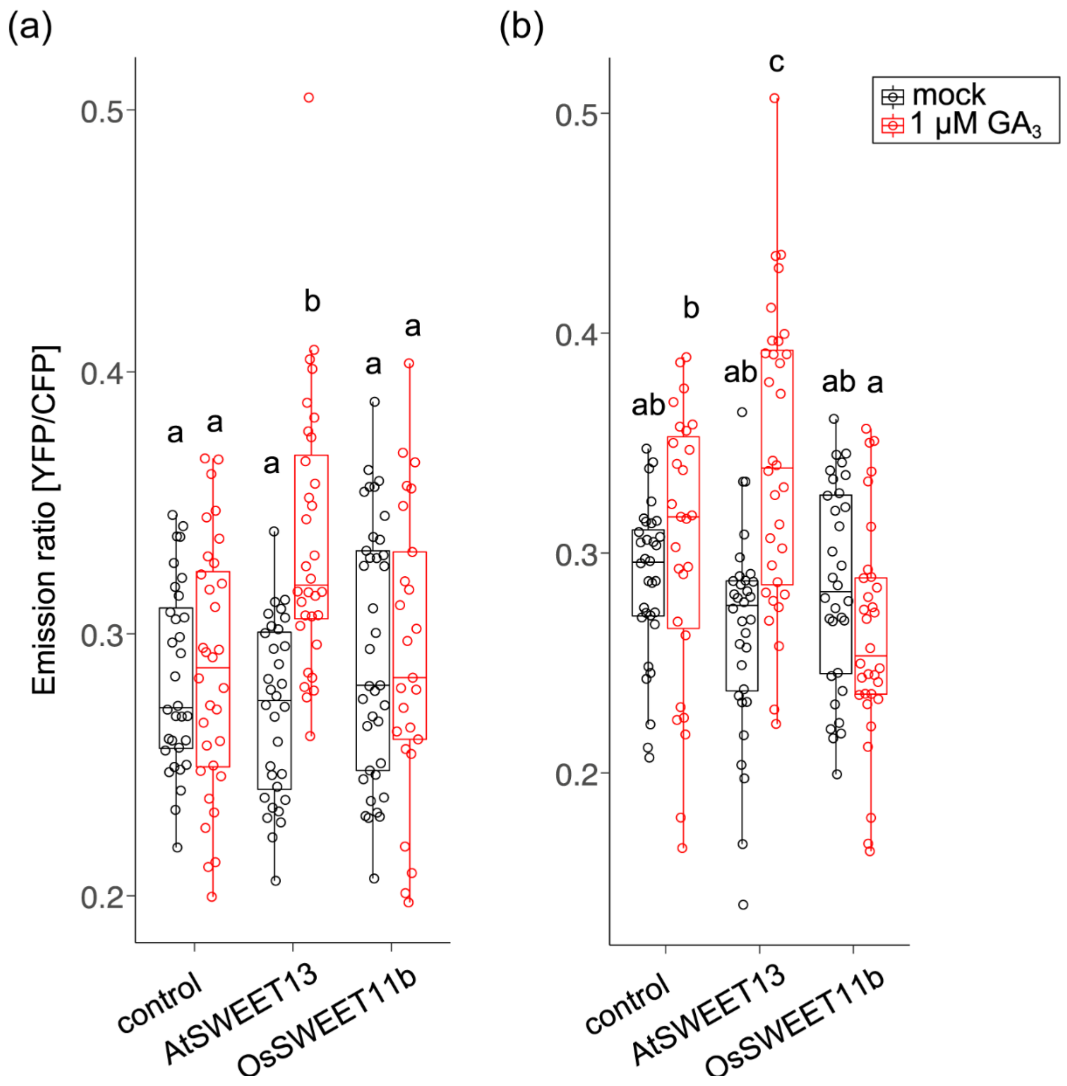


**Figure S16. $GA_3$ uptake by OsSWEET11b into human Hek293T cells.** 1 µM $GA_3$ was added for 3 h to HEK293T cells expressing the GA sensor GPS1. Empty vector and *AtSWEET13* served as negative and positive controls, respectively. Box plots show the first and third quartiles, split by the median with whiskers extending 1.5x interquartile range beyond the box. (a) Each data point represents average value of the fluorescence ratio during 3 min recordings; $n$ = 32 (empty vector, mock), 32 (empty vector, 1 µM), 32 (*AtSWEET13*, mock), 32 (*AtSWEET13* 1 µM), 37 (*OsSWEET11b*, mock), 25 (OsSWEET11b, 1 µM), and in (b); $n$ = 32 (empty vector, mock), 27 (empty vector, 1 µM), 32 (*AtSWEET13*, mock), 32 (*AtSWEET13*, 1 µM), 32 (*OsSWEET11b*, mock), 32 (*OsSWEET11b*, 1 µM)) cells. Three independent replicates were conducted. Different letters on each boxplot represents significant difference determined by One-way ANOVA with Tukey's post-hoc test ($p < 0.05$).

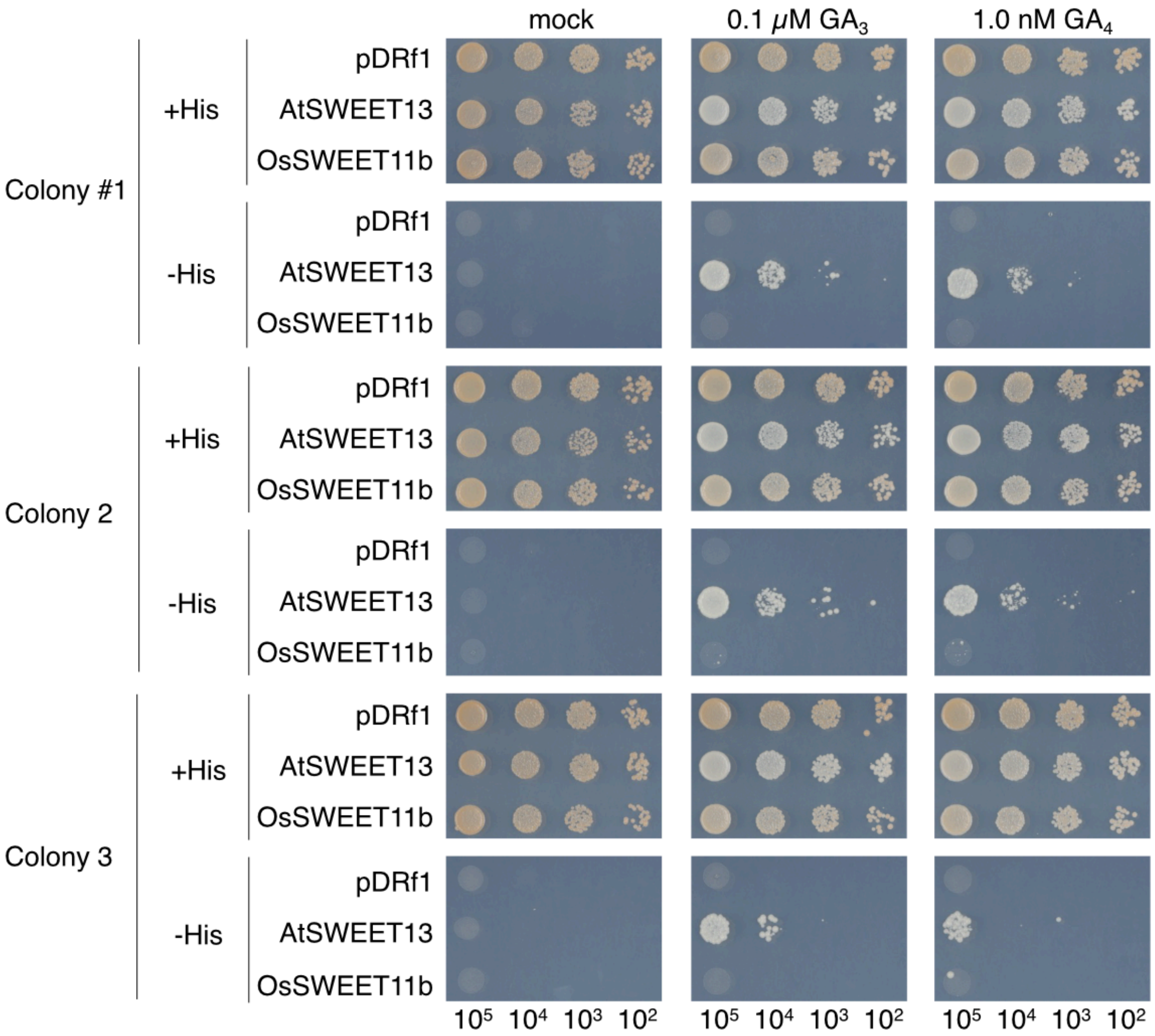


**Figure S17. Independent experiment showing $GA_3$ or $GA_4$ uptake by OsSWEET11b into yeast cells.** GA transport activity of OsSWEET11b assessed using a GA-dependent Y3H system. Yeast strain PJ69-4A carrying both pDEST22-GAI and pDEST32-GID1a and either pDRf1-OsSWEET11b, pDRf1-AtSWEET13 (positive control), or empty vector (negative control) were grown on SD(-Leu, -Trp, -Ura) or selective SD(-Leu, -Trp, -Ura, -His) medium containing 3 mM 3-AT and 0.001% (v/v) DMSO, and either 0.1 μM $GA_3$, or 1 nM $GA_4$ for 3 days at 30°C.

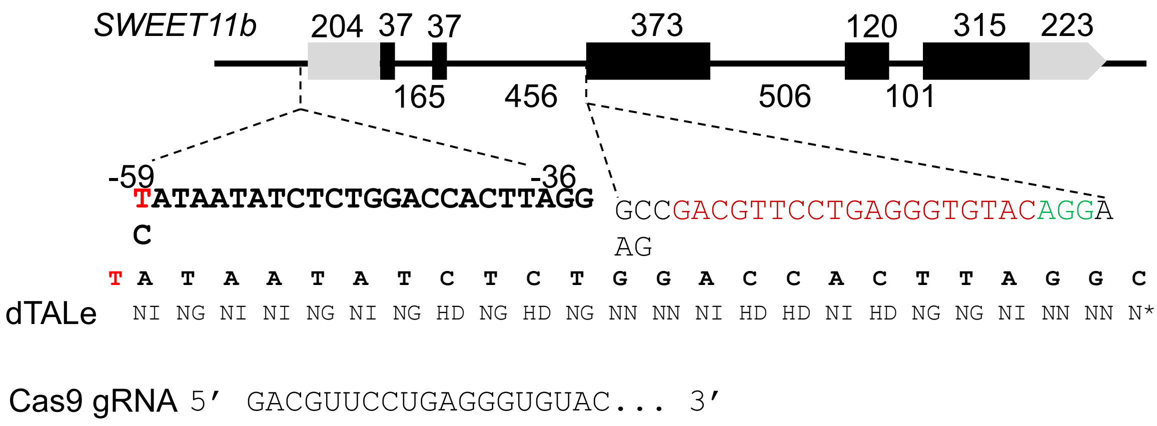


**Figure S18. Gene structure of *SWEET11b* with sequences shown for designer TALe and CRISPR/Cas9 guide RNA.** Bars represent exon and lines between present introns with DNA lengths of each component shown above (for exons) and below (for introns). The 5′- and 3′-UTRs are light grey and coding sequence in black. The sequences upstream 5′UTR are for designer TALe and sequences in exon 3 are for guide RNA with red letters for protospacer and green letter for PAM. dTALe below DNA sequences are presented by two amino acids (aa) (single letter) at positions 12 and 13 of individual 34 aa repeats. * of the last repeat indicates the $13^{th}$ aa is missing. The guide RNA (gRNA) with spacer sequence only is shown.

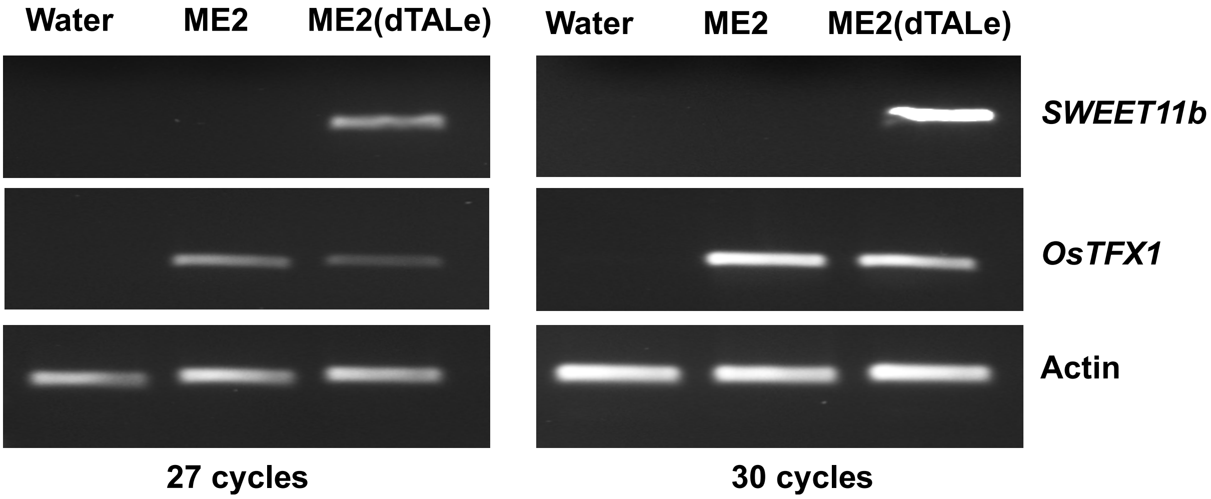


**Figure S19. Induction of *OsSWEET11b* mRNA level in rice infected with different Xoo strains.** RT-PCR of *OsSWEET11b*, the reference gene *OsTFX1* and *actin* amplified in two experiments using different cycle number. The designer TALe specifically induced *OsSWEET11b*.

### Supplementary Tables

#### Table S1 Primers used in this study

| Primer ID | Sequence (5'-3') | Purpose |
|---|---|---|
| 8NKpI_F5 | CACCGGTACCTGAGTGGTCATACGTGTCA TATTG | *ossweet11a* mutant genotyping |
| 8N3_R2 | AAGTGACTTGTGCCCATCACTTG | |
| SWT11b_F1 | GGATTGTGCCAGAGTTTTG | *ossweet11b* mutant genotyping |
| SWT11b_R1 | GTTGAGTTCAAGTTGGGGAA | |
| OsSWT11b-pcD-Bm-F | TTGGTACCGAGCTCGGCCACCATGGATC ACCTTTGGGCCTCTGTC | Mammalian expression |
| OsSWT11b-pcD-Xh-R | GGGCCCTCTAGACTCGATCAGACGGTCTT GGTGGCGCCCTTGC | |
| SWEET11bRT-F | ATGAGCGTCATCATGGTGGTG | *OsSWEET11b* expression analysis by RT-PCR |
| SWEET11bRT-R | GCTTCCGGTATTTGAAGTAG | |
| gSWEET11b-F | GTGTGACGTTCCTGAGGGTGTAC | *OsSWEET11b* CRISPR-gRNA construct |
| gSWEET11b-R | AAACGTACACCCTCAGGAACGTC | |
| SW11b_XbaI_F | GCTCTAGATCAGAAACTGGCATGTCATA | SWEET11b translational fusion GUS construct |
| SW11b_BamHI_R | CGGGATCCGACGGTCTTGGTGGCGCCCTT | |
| Ubi_F | AGAAGGAGTCCACCCTCCACC | Rice internal reference gene expression via RT-PCR |
| Ubi-R | GCATCCAGCACAGTAAAACACG | |
| SWEET11b_cds_F1 | GGGGACAAGTTTGTACAAAAAAGCAGGCTTA ATGGATCACCTTTGGGCCTC | *OsSWEET11b* coding sequence with *att*B1 and *att*B2 sites |
| SWEET11b_cds_R1 | GGGGACCACTTTGTACAAGAAAGCTGGGTAG ACGGTCTTGGTGGCGCCCT | |
| SWEET11b_cds_F2 | GGGGACAAGTTTGTACAAAAAAGCAGGCTTC ACcATGGATCACCTTTGGGCCTC | *OsSWEET11b* coding sequence cloning with *att*B1 and *att*B2 sites |
| SWEET11b_cds_R2 | GGGGACCACTTTGTACAAGAAAGCTGGGTGG ACGGTCTTGGTGGCGCCCT | |

**Table S2 Genotype and phenotype analyses of T3 plants of *ossweet11a;b* double mutants**
In frame 3-nt (-CTA) deletion introduced deletion of a single alanine at the protein level. Plants carrying 3-nt deletion or biallelic mutation in *OsSWEET11a* were not true double mutant, thus did not show male-sterility phenotype.

| Plant ID | *OsSWEET11a mutation* | *OsSWEET11b mutation* | [1]Double mutant line | Male sterility |
|---|---|---|---|---|
| Kitaake | WT | WT | | N |
| 6-4-1 | Biallelic (-C/-CTA) | +T | | N |
| 6-4-2 | -C | +T | | Y |
| 6-4-3 | Biallelic (-C/-CTA) | +T | | N |
| 6-4-4 | -C | +T | | Y |
| 6-4-5 | Biallelic (-C/-CTA) | +T | | N |
| 6-4-6 | -C | +T | | Y |
| A1-1-1 | Biallelic (-CTA/n.d) | +T | | N |
| A1-1-2 | -CTA | +T | | N |
| A1-1-3 | Biallelic (-CTA/n.d) | +T | | N |
| A1-1-4 | Biallelic (-CTA/n.d) | +T | | N |
| A1-2-1 | Biallelic (-C/-CTA) | +T | | N |
| A1-2-2 | -C | +T | | Y |
| A1-2-3 | -CTA | +T | | N |
| A1-2-4 | -C | +T | | Y |
| A1-2-5 | Biallelic (-C/-CTA) | +T | *ossweet11a-4;11b-1* | N |
| A1-2-6 | Biallelic (-C/-CTA) | +T | | N |
| A1-2-7 | Biallelic (-C/-CTA) | +T | | N |
| A1-2-8 | -CTA | +T | | N |
| A1-2-9 | -CTA | +T | | N |
| A1-2-10 | Biallelic (-C/-CTA) | +T | | N |
| 1-5-1 | -CTA | +A | | N |
| 1-5-2 | +C | +T | | Y |
| 1-5-3 | +C | +A | | Y |
| 1-5-4 | Biallelic (-CTA/+C) | Biallelic (+W) | | N |
| 1-5-5 | -CTA | Biallelic (+W) | | N |
| 1-5-6 | Biallelic (-CTA/+C) | +A | *ossweet11a-3;11b-3* | N |
| 1-5-7 | +C | Biallelic (+W) | | Y |
| 1-6-1 | -CTA | Biallelic (+W) | | N |
| 1-6-2 | +C | +A | | Y |
| 1-6-3 | +C | Biallelic (+W) | | Y |
| 1-6-4 | Biallelic (-CTA/+C) | +A | | N |
| 1-6-5 | Biallelic (-CTA/+C) | Biallelic (+W) | | N |
| 1-6-6 | Biallelic (-CTA/+C) | +T | | N |
| 1-6-7 | -CTA | Biallelic (+W) | | N |
| 1-6-8 | Biallelic (-CTA/+C) | +T | | N |
| 1-6-9 | +C | +T | | Y |

[1]Since double mutant lines were sterile, the lines used for the investigation were progenies from the plants with *OsSWEET11a* with biallelic mutations that were fertile.
